# A target capture approach for phylogenomic analyses at multiple evolutionary timescales in rosewoods (*Dalbergia* spp.) and the legume family (Fabaceae)

**DOI:** 10.1101/2021.12.07.471551

**Authors:** Simon Crameri, Simone Fior, Stefan Zoller, Alex Widmer

## Abstract

Understanding the genetic changes associated with the evolution of biological diversity is of fundamental interest to molecular ecologists. The assessment of genetic variation at hundreds or thousands of unlinked genetic loci forms a sound basis to address questions ranging from micro-to macro-evolutionary timescales, and is now possible thanks to advances in sequencing technology. Major difficulties are associated with i) the lack of genomic resources for many taxa, especially from tropical biodiversity hotspots, ii) scaling the numbers of individuals analyzed and loci sequenced, and iii) building tools for reproducible bioinformatic analyses of such datasets. To address these challenges, we developed a set of target capture probes for phylogenomic studies of the highly diverse, pantropically distributed and economically significant rosewoods (*Dalbergia* spp.), explored the performance of an overlapping probe set for target capture across the legume family (Fabaceae), and built a general-purpose bioinformatics pipeline. Phylogenomic analyses of *Dalbergia* species from Madagascar yielded highly resolved and well supported hypotheses of evolutionary relationships. Population genomic analyses identified differences between closely related species and revealed the existence of a potentially new species, suggesting that the diversity of Malagasy *Dalbergia* species has been underestimated. Analyses at the family level corroborated previous findings by the recovery of monophyletic subfamilies and many well-known clades, as well as high levels of gene tree discordance, especially near the root of the family. The new genomic and bioinformatics resources will hopefully advance systematics and ecological genetics research in legumes, and promote conservation of the highly diverse and endangered *Dalbergia* rosewoods.

## 1 INTRODUCTION

The question how biological diversity evolves is of fundamental interest in ecology and evolution, and addressing it benefits from integrative approaches (Cutter, 2013; Rissler, 2016). Investigating evolutionary processes acting at the level of populations or groups of spatially interconnected populations (metapopulations) within species typically falls within the fields of population genetics and phylogeography. By contrast, analyses of evolutionary relationships among species and patterns of diversification in higher taxonomic groups fall within the realm of phylogenetics. Though it has long been recognized that “the same ecological and evolutionary processes that cause lineage divergence can also drive speciation” (Rissler, 2016), research in these fields has traditionally relied on different conceptual approaches, analytical methods, and molecular markers, generating a false dichotomy between fields aiming to address the same underlying processes. Today, the conceptualization of common theory combined with advances in methodology leveraging on next-generation sequencing (NGS) data offer the opportunity to jointly study the processes that drive the evolution of biological diversity from micro-to macro-evolutionary timescales.

Target capture (Mamanova et al., 2010) provides an efficient approach to acquire molecular information across broad evolutionary timescales when genomic regions with varying level of diversity are included in the experimental design (Jones & Good, 2016). It requires the design of capture probes that target unique regions in the genome to prevent conflation of orthologs and paralogs, and are characterized by a conserved core for in-solution hybridization and more variable flanking regions expected to provide parsimony informative sites (Lemmon et al., 2012). Combined with high-throughput sequencing, this approach allows for the analysis of hundreds or thousands of orthologous loci in dozens to hundreds of individuals at moderate per-sample costs, and therefore strikes a good balance between locus information content and scalability to high numbers of individuals, including museum specimens (de La Harpe et al., 2017; Brewer et al., 2019). Hence, target capture holds a great potential to bridge the divide between phylogenetics, phylogeography and population genetics (de La Harpe et al., 2017; Nicholls et al., 2015; Rissler, 2016) and has increasingly been applied at macro-evolutionary, phylogeographic and micro-evolutionary timescales in a wide range of animals (e.g., Faircloth et al., 2012; Lemmon et al., 2012; Prum et al., 2015) and plants (e.g., de La Harpe et al., 2018; Koenen et al., 2020a; Mandel et al., 2014).

A global probe set targeting 353 putatively single-copy protein-coding genes has recently been developed for flowering plants (Angiosperms353; Johnson et al., 2019). Recent studies in various plant families have shown that the Angiosperms353 probe set represents a cost-effective resource to resolve phylogenetic relationships at the level of plant orders (e.g., Thomas et al., 2021), families (e.g., Siniscalchi et al., 2021), or at the infrageneric level (e.g., Ottenlips et al., 2021). However, several comparisons revealed that micro-evolutionary relationships are often better resolved when targeting more loci using taxon-specific probe sets (e.g., Shah et al., 2021; Siniscalchi et al., 2021; Ufimov et al., 2021). The development of taxon-specific probe sets therefore remains valuable for detailed phylogenetic and population genetic analyses (Yardeni et al., 2021).

Beside challenges associated with the *de-novo* probe design, processing and analysis of high-throughput sequencing data often involves complex and computationally demanding calculations. Target capture data are often analyzed using the PHYLUCE (Faircloth, 2016) or HYBPIPER (Johnson et al., 2016) bioinformatic pipelines. PHYLUCE was developed for analysis of sequences flanking ultraconserved genomic elements and has mainly been used at macro-evolutionary and phylogeographic timescales in animal systems, whereas HYBPIPER is optimized for datasets derived from probes designed in exons using HYB-SEQ (Weitemier et al., 2014). There is thus a need for existing tools to be expanded with pipelines that are applicable at deep to shallow evolutionary timescales (de La Harpe et al., 2017), while being independent from high-quality annotated genomes or transcriptomes.

*Dalbergia* L.f. (Fabaceae) is a pantropical and ecologically diverse plant genus with c. 270 currently accepted species (WCVP, 2021), some of which have been described relatively recently (e.g, Adema et al., 2016; Lachenaud, 2016; Wilding et al., 2021a, 2021b). Numerous arborescent species are a source of rosewood (Bosser & Rabevohitra, 2002; Prain, 1904), a high-quality timber sought-after on the international market and cause of conservation concern (Schuurman & Lowry, 2009; Waeber et al., 2019). National and international regulations have been established, aiming at sustainable exploitation and revenues (Barrett et al., 2013; CITES, 2020), but illegal logging and trade continues (UNODC, 2016b, 2020; Vardeman & Runk, 2020). The effective implementation of regulations demands that species are reliably recognized and that extant population sizes are estimated to assess the potential threat status. Developing a comprehensive understanding of species diversity in *Dalbergia* and their evolutionary history, as well as a thorough knowledge of the ecology and distribution of many traded species, has been hampered by several factors. There is a shortage of collections and experts focusing on this taxonomically challenging genus, and current treatments heavily rely on leaves and flowers and/or fruits for identification (Bosser & Rabevohitra, 2002; de Carvalho, 1997; Lachenaud, 2016), which are rarely encountered together in the field. As a result, the taxonomy of the genus is in need of extensive revision (Wilding et al., 2021a), which could be supported by phylogenomic analyses targeting the nuclear genome (Crameri, 2020).

Motivated by the need for genomic resources to inform a reliable taxonomy and foster conservation practice, we introduce a target capture approach for anchored phylogenomic analyses in *Dalbergia* (Dalbergia2396 set). This genus belongs to the third largest angiosperm family (Fabaceae, a.k.a. Leguminosae or legume family), which is subject to extensive research in areas such as systematics (LPWG, 2017), ecology (Sprent et al., 2017), evolution (Koenen et al., 2021), speciation and rapid radiations (Hughes & Eastwood, 2006), and contains many agricultural crops (Mousavi-Derazmahalleh et al., 2018; Zhuang et al., 2019). This motivated us to further explore the applicability of our approach for analyses across the entire legume family, which resulted in a second probe set (Fabaceae1005 set). Both probe sets represent a subset of 6,555 conserved target regions distributed across the nuclear genome, derived from a combination of divergent reference capture using five published legume genomes, and a *de novo* assembly of a *Dalbergia* transcriptome. We also introduce a dedicated bioinformatics pipeline named CAPTUREAL supporting the analysis of high-throughput target capture sequencing data, with special emphasis on streamlined applicability, parallelization, and graphical output for informed parameter choices. The pipeline is designed for general application to target capture datasets, modular, and therefore easily customizable. We demonstrate the application of our approach to resolve phylogenetic relationships in the economically important and conservation-relevant genus *Dalbergia*. We then explore the utility for phylogenomic analyses at much deeper timescales by analyzing target capture data of various legume subfamilies. Finally, we test the utility of this approach at a micro-evolutionary scale, and assess genetic variation among individuals and populations of two closely related *Dalbergia* species from Madagascar.

## 2 MATERIALS AND METHODS

### 2.1 Design of target capture probes and reference sequences

We produced a transcriptome assembly of a cultivated individual of *Dalbergia madagascariensis* subsp. *antongilensis* Bosser & R. Rabev., based on 63 million paired-end sequencing reads generated on an Illumina® HiSeq^TM^ 2000 platform. We performed *de novo* assembly of the transcriptome using Trinity release 2012-01-25 (Grabherr et al., 2011), resulting in 146,484 scaffolds, which were between 201 and 17,129 bp long, with a mean length of 815 bp (see Supplementary Methods). We then pairwise aligned the *Dalbergia* transcriptome with reference genomes of five legume species available in public databases to generate a set of 12,049 probes from 6,555 conserved target regions (see Supplementary Methods). This probe set was used for synthesis of hybridizing probes at myBaits® Custom Target Capture Kits (Arbor Biosciences; https://arborbiosci.com).

### 2.2 Taxon sampling for target capture probes validation

We created three taxon sets with contrasting levels of evolutionary divergence, ranging from subfamilies to species to populations. The subfamily set (Table S1) included five of the six legume subfamilies, as recognized in the most recent treatment (LPWG, 2017), and comprised 104 individuals (110 samples, six replicates; 99 species including three outgroups). Three species of *Polygala* Tourn. ex L. (Polygalaceae) were included as the outgroup for the subfamily set. The species set (Table S2) included members of the closely related genera *Dalbergia* (at least 19 species)*, Machaerium* Pers. (three species) and *Ctenodon* Baill. sensu Cardoso et al. (2020) (two species) and comprised 60 individuals (63 samples, three replicates; at least 26 species including two outgroups). Two species of *Aeschynomene* L. sensu stricto (s.str.) sensu Cardoso et al. (2020) were included as the outgroup for the species set. The population set (Table S3, Figure S4) included 51 individuals in total, 29 attributed to *D. monticola* Bosser & R. Rabev. from four sampling locations, and 22 attributed to *D. orientalis* Bosser & R. Rabev. from eleven sampling locations.

### 2.3 Library preparation, target capture and sequencing

We extracted total genomic DNA from silica gel dried leaf tissue (185 extractions) or museum specimens deposited at the Paris (P) herbarium (11 extractions) using the CTAB protocol (Doyle and Doyle, 1987) or the DNeasy® Plant Mini Kit (Qiagen). We quantified DNA using the QuantiFluor® dsDNA system for a Quantus^TM^ fluorometer (Promega) and checked DNA integrity on 1.5% agarose gels for a subset of samples. We prepared genomic DNA libraries for each sample using the NEBNext® Ultra II DNA Library Prep Kit for Illumina® (New England Biolabs), following manufacturer’s instructions. We individually indexed samples to be pooled within the same sequencing lane during the PCR enrichment step using NEBNext® Multiplex Oligos for Illumina® (single-indexed with E7335 and E7500 kits, or dual-indexed with E6440 kit, New England Biolabs). We performed in-solution hybridization and target enrichment using our 12,049 tiled RNA probes. We pooled up to six individually indexed libraries during the hybridization step using a stratified random assignment of libraries to hybridization reactions. Stratification aimed at optimizing the sequencing coverage across samples and consisted in avoiding pooling of close relatives of *Cajanus cajan* with more distantly related samples, and of museum specimens with silica gel dried leaf material. We obtained short read data by combining sequencing runs from an Illumina® MiSeq^TM^ (2×300 bp paired-end sequencing, 99 libraries) at the Genetic Diversity Centre (GDC) Zurich, an Illumina® HiSeq^TM^ 4000 (2×150 bp paired-end sequencing, 88 libraries) at the Functional Genomics Center Zurich (FGCZ) or Fasteris SA (Plan-les-Ouates, Switzerland), and an Illumina® NovaSeq^TM^ 6000 SP flow cell (2×150 bp paired-end sequencing, 9 libraries) at the FGCZ. We repeated DNA extraction, hybridization and target enrichment sequencing for nine individuals (replicates) to assess reproducibility. One sample (*Hassold 565*) was represented in each taxon set, nine samples were represented in both the species and population sets, and nineteen samples were represented in both the subfamily and species sets.

### 2.4 CAPTUREAL bioinformatics pipeline

The bioinformatic pipeline CAPTUREAL was developed for this project and is accessible on Github (https://github.com/scrameri/CaptureAl) as a documented sequence of scripts. These include bash and R scripts (R Core Team, 2020) to manage and visualize data with APE version 5.3 (Paradis & Schliep, 2018), DATA.TABLE version 1.12 (Dowle & Srinivasan, 2019), and TIDYVERSE version 1.3.0 (Wickham et al., 2019). Where appropriate, computations are carried out for multiple samples or target regions in parallel using GNU PARALLEL (Tange, 2011). The CAPTUREAL pipeline streamlines the mapping of quality-trimmed reads to target regions, the exclusion of loci targeting multi-copy genes and taxa with insufficient data coverage, and the alignment of orthologous loci for downstream phylogenetic analyses. At various critical steps, the pipeline outputs summary statistics and graphs that inform the user on the effects of specific filtering parameters, allowing for informed parameter choices.

The pipeline is divided into seven steps to process quality-filtered reads. Steps 1 to 5 are always required, and 1) map the sequencing reads to target regions, 2) assemble mapped reads separately for each target region, 3) identify the most-likely orthologous contigs, 4) identify taxa and target regions with high capture sensitivity and specificity, and 5) create trimmed alignments of the kept taxa and target regions. Steps 6 and 7 are optional, and 6) combine physically neighboring and overlapping alignments to 7) generate longer and more representative reference sequences as starting points for reiteration of steps 1 to 5. Such reiteration can improve mapping success, and mitigates potential biases arising from the reference sequences used (Hahn et al., 2013).

In our analyses, we executed the pipeline separately and iteratively for different taxon sets. We first applied steps 1 to 5 to twelve representative samples each from the subfamily and species sets, followed by steps 6 and 7 to generate longer and taxon-specific reference sequences for target regions that can each be efficiently recovered in these taxon sets, and then reiterated steps 1 to 5 for all samples of the subfamily and species sets using the new reference sequences and more stringent target region filtering parameters (see Tables S1–S3 and Supplementary Methods for details). We also performed steps 6 and 7 after the second iteration of the species set analysis to produce reference sequences for analysis of the population set. Bioinformatic analyses were carried out on a multi-core LINUX server (GDC Zurich) or on the EULER scientific compute cluster (ETH Zurich). The sequence of executed commands and the chosen parameters are provided in Supplementary Methods.

#### 2.4.1 Step 1: Read mapping

Quality-filtered reads of each sample are mapped against the reference sequences (one sequence per target region) using the BWA-MEM algorithm (Li, 2013). The minimum alignment score and mapping quality can be adjusted as needed. Coverage statistics are computed using SAMTOOLS (Li & Durbin, 2009) and BEDTOOLS (Quinlan & Hall, 2010), and target regions are visually filtered for adequate coverage across samples using *filter.visual.coverages.R*, which allows to apply filtering thresholds that are informed by visualizations of coverage statistics (see Supplementary Methods). The main output of step 1 are BAM files a list of retained target regions.

#### 2.4.2 Step 2: Sequence assembly

Read pairs are extracted from quality-filtered reads when at least one read mapped to any of the retained target regions with the specified minimum mapping quality. Extracted reads are assembled separately for each sample and region using DIPSPADES (Safonova et al., 2015) based on haplocontigs generated by SPADES (Bankevich et al., 2012, see Supplementary Methods). The main output of step 2 are consensus contiguous sequences (contigs hereafter) for each sample and each target region.

#### 2.4.3 Step 3: Orthology assessment

Sequence assembly may yield multiple contigs per sample for some target regions, e.g., due to capture of several fragments of the same genomic region (e.g., in degraded museum specimens), or due to unspecific capture of paralogs (Johnson et al., 2016). The most likely orthologous contig(s) of each sample in each target region are determined using an exhaustive Smith-Waterman alignment (Smith & Waterman, 1981) between all contigs and the reference sequences using EXONERATE (Slater & Birney, 2005). The best-matching contig is defined based on the EXONERATE alignment statistics as the most likely orthologous contig for each sample and target region, and further contigs that did not overlap with one another or the best-matching contig, but aligned with a sufficient alignment score to other parts of the target region are retained. These contigs likely represent fragments of the same region, and can therefore be combined with the best-matching contig to form a contiguous sequence (orthologous contig hereafter, see Supplementary Methods). The main output of step 3 is a single-sequence FASTA file with the putative orthologous contig for each sample and each target region.

#### 2.4.4 Step 4: Sample and region filtering

Successful target capture depends on whether sequence data can be collected for a high proportion of target regions (capture sensitivity, Jones & Good, 2016) in a high proportion of focal taxa, and whether the captured sequences are orthologs of the target regions (capture specificity). Target regions are visually filtered for high capture sensitivity and specificity across focal taxa using *filter.visual.assemblies.R*, which allows to apply filtering thresholds that are informed by visualizations of EXONERATE alignment statistics generated in step 3. These thresholds can be set globally to remove generally poorly sequenced samples or target regions, but they can also be set as the minimum fraction of samples required to pass a specified filtering threshold in order for a target region to be retained. Taxon groups can be defined, in which case the required capture sensitivity and specificity parameters need to be met in all considered taxon groups separately, thus preventing target regions from being poorly represented in rare taxon groups (see Supplementary Methods). The main output of step 4 is a list of samples and a list of target regions to keep.

In our analyses, we defined the four subfamilies represented by multiple taxa as taxon groups in the subfamily set. In the species set we defined four taxon groups based on our preliminary phylogenetic results and phylogenetic relationships inferred by Hassold et al. (2016). These were subgroup (SG) 1 (species with large flowers and paniculate inflorescences), SG2 (species with large flowers and racemose inflorescences), SG3 (species with small flowers from East Madagascar), and SG4 (species with small flowers from West and North Madagascar).

#### 2.4.5 Step 5: Target region alignment and alignment trimming

A multi-sequence FASTA file is generated for all retained target regions, containing the respective orthologous contigs of all retained samples. Sequences are then aligned using MAFFT (Katoh & Standley, 2013), allowing for different alignment options. Alignments are trimmed at both ends until an alignment site shows nucleotides across a specified minimum fraction of aligned sequences, along with a specified maximum nucleotide diversity (i.e., the mean number of base differences between all sequence pairs). In addition, internal trimming is performed by only keeping sites with nucleotides in a specified minimum fraction of aligned sequences. Potential mis-assemblies or mis-alignments at contig ends are further resolved using a sliding window approach that identifies and masks sequences with large deviations from the alignment consensus (see Supplementary Methods). The main output of step 5 are potentially overlapping trimmed alignments for each kept target region.

#### 2.4.6 Step 6: Merging of overlapping alignments

Shorter but physically close target regions facilitate sequence assembly in lower-quality samples but can lead to overlaps in trimmed alignments of neighboring target regions. Such overlaps can be identified by aligning consensus sequences of target region alignments. Specifically, consensus sequences are generated by calling IUPAC ambiguity codes if a given minor allele frequency threshold is reached, or a gap if a given base frequency threshold is not reached. Local alignments between different consensus sequences are identified using BLAST+ version 2.7.1 (Camacho et al., 2009), and filtered for non-reciprocal hits between alignment ends of target regions located on the same linkage group. Orthologous contigs that are part of different, overlapping alignments are then aligned using MAFFT. The resulting merged alignments are then collapsed to represent different orthologous contigs of the same individual as a single sequence, a process that can be visually inspected if needed. Trimming is then applied as in step 5, and sets of two to several consecutively overlapping alignments are then each replaced by a single merged alignment if merging was successful (see Supplementary Methods). The main output of step 6 are non-overlapping trimmed alignments for each kept target region.

#### 2.4.7 Step 7: Generation of representative reference sequences

To mitigate potential biases arising from the reference sequences used, a new set of target region reference sequences can be generated based on the target region alignments generated in the two previous steps. For this purpose, a consensus sequence is generated for each alignment as in step 6, but separate consensus sequences can be generated for different specified taxon groups (see step 4). These sets of taxon group specific consensus sequences are then aligned, and representative consensus sequences are generated as in step 6 (see Supplementary Methods). These taxon-specific reference sequences are the main output of step 7 and can be used to refine mapping, assembly and alignment by reiterating steps 1 to 5.

#### 2.4.8 Alignment assessment and filtering

We characterized all non-overlapping trimmed alignments for the number of gaps, gap ratio (i.e, the fraction of non-nucleotides in the alignment), total nucleotide diversity, average nucleotide diversity per site, and alignment length, as well as the number and proportion of segregating and parsimony informative sites. We then visually filtered alignments using *filter.visual.alignments.R*, which allows to apply filtering thresholds that are informed by visualizations of alignment statistics (see Supplementary Methods). We used the filtered alignments after the second iteration of step 6 for phylogenetic analyses.

### 2.5 Phylogenetic analyses

We performed phylogenetic analyses with both the subfamily and species sets, using a supermatrix (concatenation) approach and a gene tree summary approach. For the supermatrix approach, we ran maximum likelihood searches on the concatenated alignments using RAXML version 8.2.11 (Stamatakis, 2014) with rapid bootstrap analysis and search for the best-scoring tree in the same run (-f a option), 100 bootstrap replicates, and the GTRCAT approximation of rate heterogeneity (see Supplementary Methods). For the gene tree summary approach, we ran RAXML jobs separately for each alignment using the same settings as for the supermatrix approach to generate gene trees. Following Zhang et al. (2018), we collapsed branches in gene trees if they had bootstrap support values below 10 using NEWICK utilities (Junier & Zdobnov, 2010), and we performed species tree analyses with ASTRAL-III version 5.6.3 (Mirarab et al., 2014; Zhang et al., 2018) and standard parameters, except for full branch annotation (see Supplementary Methods). For the subfamily set, we additionally evaluated the quartet support for fifteen different subfamily topologies (i.e., all possible topologies with Caesalpinioideae, Dialioideae, Papilionoideae and (Cercidoideae, Detarioideae) as ingroups; Figure S2), using the tree scoring option in ASTRAL-III and a file with the assignment of taxa to subfamilies or the outgroup. All phylogenetic trees were displayed using GGTREE version 2.0.2 (Yu et al., 2016).

### 2.6 Population genetic analyses

We carried out population genetic analyses for the population dataset. We mapped quality-filtered reads against the target region reference sequences that were representative of the species set after the second iteration using BWA-MEM. We verified efficient recovery of target regions by plotting heatmaps of coverage statistics, removed PCR duplicates using PICARD TOOLS version 2.21.3 (Broad Institute, 2019), and capped excessive coverage to 500 using *biostar154220.jar* (Lindenbaum, 2015). We then called variants using FREEBAYES version 1.1.0-3-g961e5f3 (Garrison & Marth, 2012) and standard parameters, except for a minimum alternate fraction of 0.05, a minimum repeat entropy of 1, and evaluation of only the four best alleles. Variants were filtered using VCFTOOLS version 0.1.15 (Danecek et al., 2011) and VCFLIB version 1.0.1 (Garrison, 2012), which was also used to decompose complex variants (see Supplementary Methods). We then used VCFR version 1.10.0 (Knaus & Grünwald, 2017) and ADEGENET version 2.1.1 (Jombart, 2008; Jombart & Ahmed, 2011) to generate *genind* and *genlight* objects that represented the SNP allele table with associated metadata such as individual missingness, species identification, and sampling location. We used the SNP subset with zero missingness to conduct principal component analysis (PCA) based on the centered covariance matrix, as well as to calculate a neighbor-joining (NJ) tree (Saitou & Nei, 1987) on Nei’s genetic distances, as implemented in POPPR version 2.8.1 (Kamvar et al., 2014). We also used the allele table to create a SNP subset for population clustering analysis using STRUCTURE version 2.3.4 (Pritchard et al., 2000). Specifically, we kept SNPs with genotype data in at least 95% of individuals, and we randomly sampled up to three SNPs per target region for linkage disequilibrium pruning and computational ease. STRUCTURE analyses were performed for one to ten demes (*K*), using 110,000 iterations, including a burn-in period of 10,000 iterations, with ten replicates per simulation (see Supplementary Methods). Replicate STRUCTURE results were aligned and visualized using CLUMPAK (Kopelman et al., 2015) and default settings.

## 3 RESULTS

### 3.1 Two probe sets for target capture across legumes and *Dalbergia*

We obtained 0.13 to 13.76 (median: 1.56) million raw read pairs per sample, of which we retained 86.55% to 99.34% (median: 93.82%) after quality trimming (Tables S1 – S3). In the first iteration applied to twelve representative samples, reads mapped to 6,519 or 6,287 of the 6,555 initial target regions in the subfamily or species set, respectively (step 1). Of these we retained 3,436 or 4,908 target regions, which showed adequate coverage across taxon groups. After assembly (step 2) and orthology assessment (step 3), 2,710 or 4,181 target regions passed the region specificity and sensitivity filters of lower stringency (step 4). Following alignment and trimming (step 5), overlapping portions in 207 or 377 regions were successfully merged, resulting in 2,468 or 3,736 non-overlapping trimmed alignments (step 6). Longer and more representative consensus sequences were generated from these target regions (step 7) and used as references for mapping quality-trimmed reads of the complete taxon sets (step 1, see Tables S1 and S2). We retained 1,917 or 3,418 target regions with adequate coverage (Figures S5 and S6), of which 1,020 or 2,407 passed the specificity and sensitivity filters of higher stringency (step 4) after assembly. Merging of overlapping alignments in 15 or 11 regions yielded 1,005 (subfamily set) or 2,396 (species set) distinct alignments (step 5), of which 666 represented the same regions in both sets. The corresponding Fabaceae1005 and Dalbergia2396 probe sets, along with refined taxon-specific reference sequences are deposited on Dryad. Corresponding gene annotations in the *Cajanus cajan* genome are given in Tables S4 and S5. For phylogenetic analyses, we excluded 19 or 7 alignments with a gap ratio above 0.35 or 0.3 or a nucleotide diversity above 0.35 or 0.15, leaving 986 (subfamily set) or 2,389 (species set) alignments.

Quality-trimmed reads mapped to all 2,396 target regions in the population set (step 1) using reference sequences that were representative of the species set after the second iteration for mapping (Figure S7). Variant calling resulted in `203,916 raw variants and` 116,500 filtered SNPs after decomposing complex variants, of which 60,204 (51.68%) were bi-allelic with no missing data and were used for PCA and NJ tree reconstruction. Random sampling of up to three SNPs per target region resulted in a subset of 5,042 SNPs for STRUCTURE analyses.

### 3.2 Phylogenomic analyses across legumes

Phylogenetic analysis of 986 alignments recovered each of the five sampled subfamilies as monophyletic, and many well-established clades and relationships received ≥95% support using both the gene tree summary method ASTRAL-III (Figure 1) and the supermatrix method (Figure S1). These included the subfamilies Cercidoideae and Detarioideae found to be sister taxa, the mimosoid clade within the recently re-circumscribed subfamily Caesalpinioideae (LPWG, 2017), as well as the Angylocalyceae-Dipterygeae-Amburaneae (ADA, Cardoso et al., 2012), Cladrastis (Wojciechowski, 2013) and Meso-Papilionoideae (Wojciechowski, 2013) clades within Papilionoideae. We also recovered the Sophoreae and Genisteae clades (Cardoso et al., 2013) within Genistoids sensu lato (s.l.) (Cardoso et al., 2012; Wojciechowski et al., 2004). Within the Dalbergioids s.l. (Wojciechowski et al., 2004), we recovered the Amorpheae clade (McMahon & Hufford, 2004) as sister to the rest of the group, which includes the Dalbergioids s.str. clade (Lavin et al., 2001), containing the *Adesmia*, *Pterocarpus* and *Dalbergia* subclades (Lavin et al., 2001), respectively. *Ctenodon brasilianus* (Poir.) D.B.O.S.Cardoso, P.L.R.Moraes & H.C.Lima and *C. nicaraguensis* (Oerst.) A.Delgado were found to be more closely related to *Machaerium* than to *Aeschynomene*. Within the Non-Protein-Amino-Acid-Accumulating (NPAAA) clade (Cardoso et al., 2012; Wojciechowski et al., 2004), we recovered the Millettioid s.l. clade (Wojciechowski et al., 2004), containing the genera *Indigofera* and *Millettia*, and the Phaseoleae s.l. (Vatanparast et al., 2018), as well as the Hologalegina (Wojciechowski, 2013) clade, including the Robinioids and the inverted-repeat-lacking clade (IRLC, Wojciechowski et al., 2004).

**FIGURE 1.**
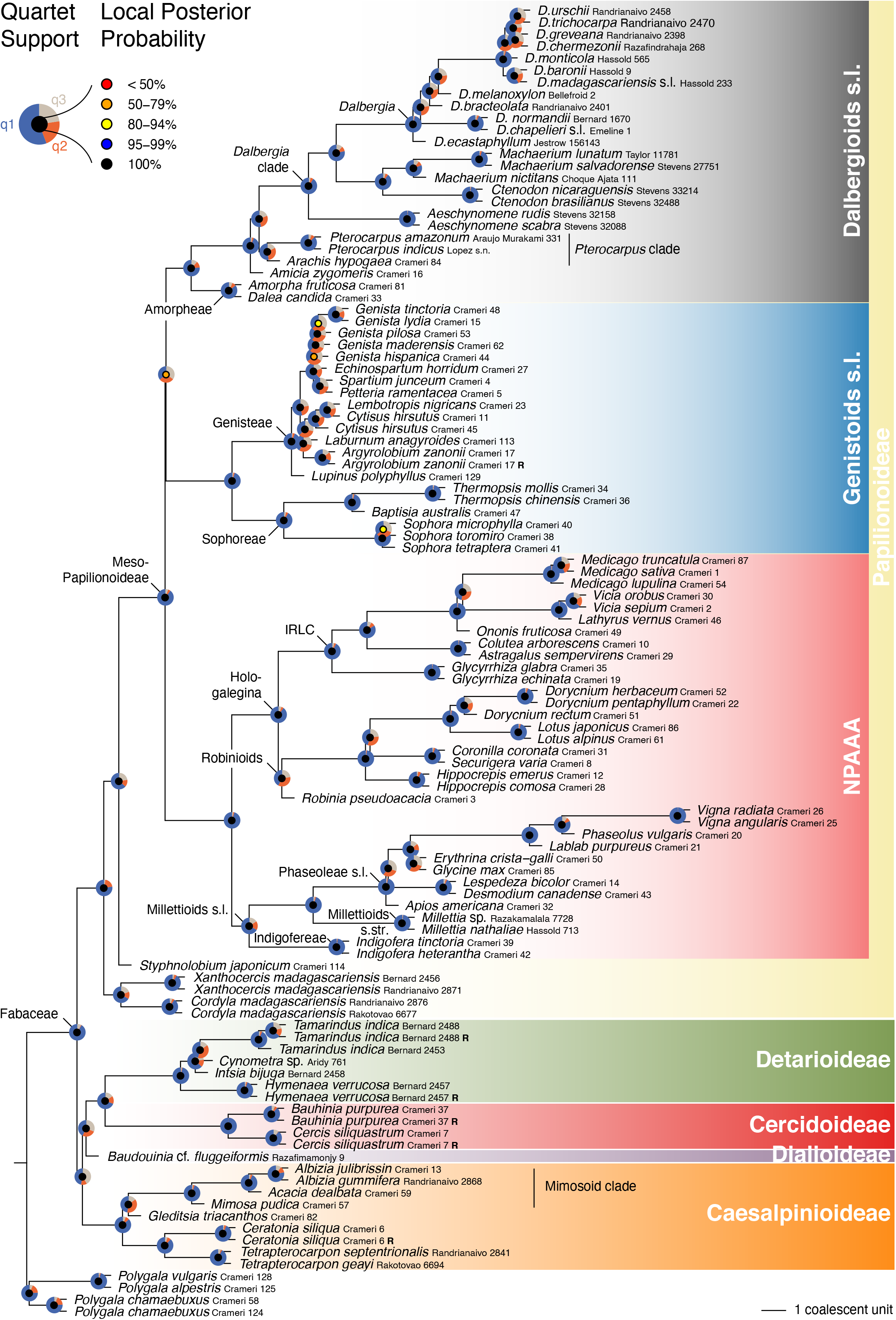
Coalescent-based phylogeny of the subfamily set (n = 110) inferred using ASTRAL-III on 986 gene trees. Pie charts on each node denote the fraction of gene trees that are consistent with the shown topology (q1; blue), and with the first (q2; orange) and second (q3; gray) alternative topologies. Local posterior probabilities are shown as color-coded circles on each node (see inset legend). Replicate specimens are labelled with a bold ‘R’. 860 gene trees (87.22%) had missing taxa. The overall normalized quartet score was 88.82%.

Other relationships among subfamilies remained unresolved using both phylogenetic methods (Figure 1, Figure S1). In particular, a clade comprising Caesalpinioideae, Cercidoideae, Detarioideae and Dialioideae as sister group to Papilionoideae was not supported in the supermatrix tree, and was recovered in only 47% of quartet trees. We evaluated quartet scores (i.e., the fraction of induced quartet trees) of fourteen further topologies for relationships among sampled subfamilies (Figure S2) using the tree scoring option in ASTRAL-III in combination with a file that mapped taxa to subfamilies or to the outgroup. The subfamily topology presented in Figure 1 showed the highest normalized quartet score (38.40%). Two alternative topologies received a similar normalized quartet score of 38.36% (Figure S2) and involved a clade composed of Caesalpinioideae and Papilionoideae. Further contentious relationships between major groups concerned the three clades within Meso-Papilionoideae, where the clade formed by Dalbergioids s.l. and Genistoids s.l. was recovered only in 36% of quartet trees, and in relationships within Caesalpinioideae, Detarioideae, and Genisteae. All except one genus with multiple sampled accessions were recovered as monophyletic, the exception being *Cytisus*, which was paraphyletic with respect to *Lembotropis nigricans*. Pairs of replicates each grouped together (Figure 1, Figure S1).

### 3.3 Phylogenomic analyses in *Dalbergia*

Phylogenetic analysis of 2,389 alignments recovered samples of *Dalbergia* as monophyletic with ≥95% support using both ASTRAL-III (Figure 2) and the supermatrix method (Figure S3). Within *Dalbergia*, we recovered two large and exclusively Malagasy clades, which we name Madagascar Supergroup I and II. All Malagasy species represented by multiple accessions were recovered as highly supported clades, with the exception of *D. normandii*. Four non-Malagasy *Dalbergia* specimens and *D. bracteolata* Baker were each found to represent separate lineages.

**FIGURE 2.**
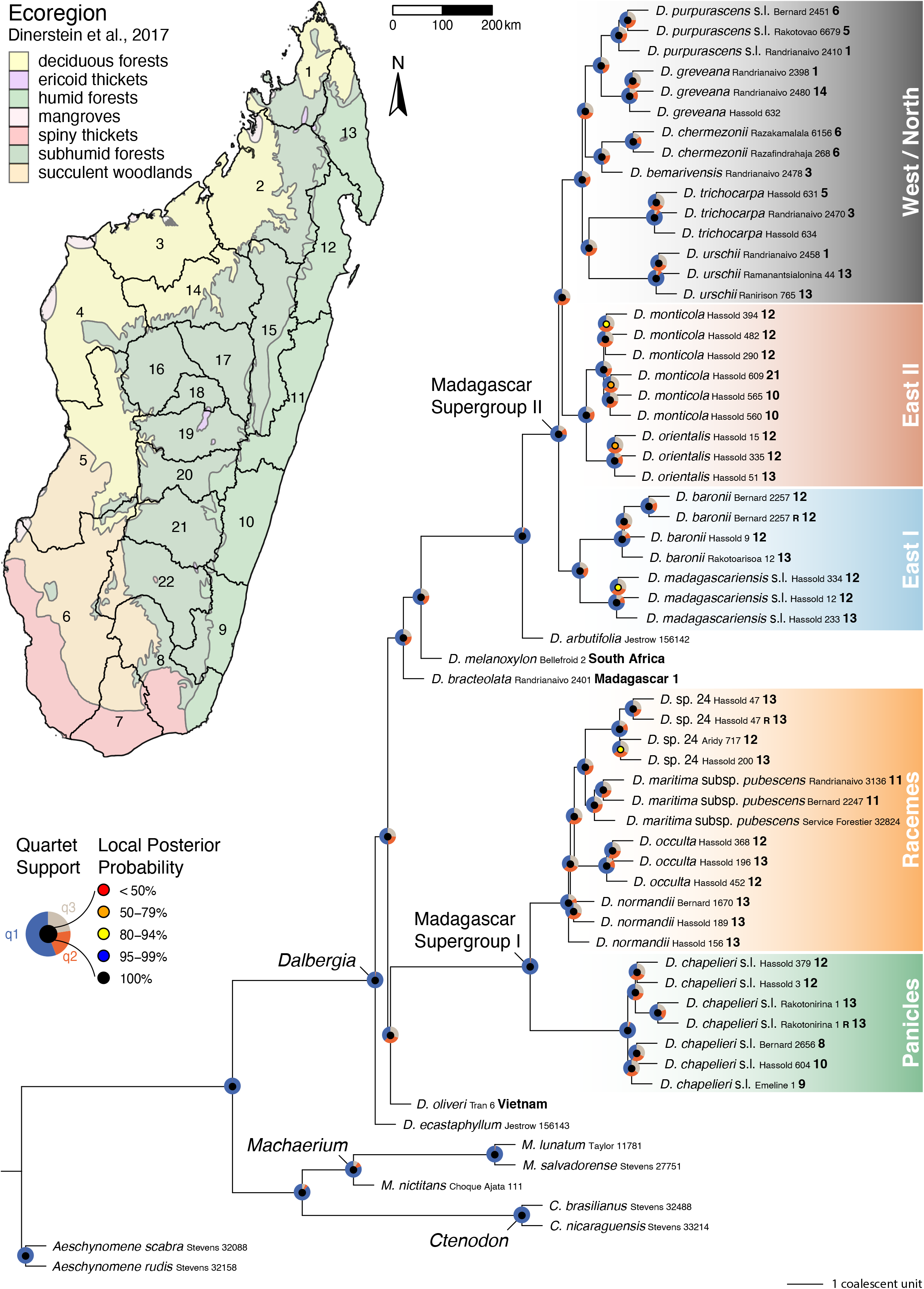
Coalescent-based phylogeny of the species set (n = 63) inferred using ASTRAL-III on 2,389 gene trees. Pie charts on each node denote the fraction of gene trees that are consistent with the shown topology (q1; blue), and with the first (q2; orange) and second (q3; gray) alternative topologies. Local posterior probabilities are shown as color-coded circles on each node (see inset legend). The geographic origins of accessions from Madagascar are indicated as bold numbers in the tree, which correspond to political regions of Madagascar, as well as to ecological regions (see inset map). Replicate specimens are labelled with a bold ‘R’. 1,014 gene trees (42.44%) had missing taxa. The overall normalized quartet score was 85.42%.

Within Supergroup I, one clade comprised samples of *Dalbergia chapelieri* s.l., while the remaining samples belonged to a sister group containing three monophyletic species and a basal and paraphyletic *D. normandii*. Within Supergroup II, two clades contained species distributed in the humid east of Madagascar, while the third contained species distributed in the seasonally dry west and north of the island. Within *D. chapelieri* s.l. and *D. monticola*, which were each represented by six individuals, we observed geographic structure, with specimens from northeast and southeast Madagascar forming sister groups. Pairs of replicates each grouped together (Figure 2, Figure S3).

### 3.4 Population genomic analyses

Principal component analysis revealed three distinct clusters of individuals along principal component (PC) 1 (explaining 27.58% of the total variation) and PC 2 (11.26%; Figure 3). Individuals of *D. orientalis* separated along PC1, while individuals originally attributed to *D. monticola* formed two distinct groups mainly along PC2. The unexpected smaller cluster (in purple) comprised samples from a single broad sampling location in north-eastern Madagascar (location 5, see Figure S4 and Table S3) where both *D. monticola* and *D. orientalis* were also collected. The same three clusters were also recovered in STRUCTURE analyses (Figure S8), where biologically meaningful clustering solutions were found for *K* = 2 (separating *D. orientalis* from the rest) and *K* = 3 (further separating the unexpected smaller cluster; Figure S9). Within *D. orientalis* and the larger cluster of presumed *D. monticola*, the NJ tree reflects isolation by distance at a broad geographical scale, separating specimens from north-eastern (locations 1 to 6), central-eastern (locations 7 and 8) and south-eastern Madagascar (locations 9 to 13; Figure 3). A similar geographic pattern was recovered by STRUCTURE assuming *K* = 5 (Figure S8), although that clustering solution received much lower support (Figure S9). Clustering solutions assuming higher *K* did not recover additional meaningful structure. *K* = 7 showed an unrealistic probability by *K* of 1 (Figure S9B), which may be related to the presence of ‘ghost clusters’ with near-zero admixture proportions.

**FIGURE 3.**
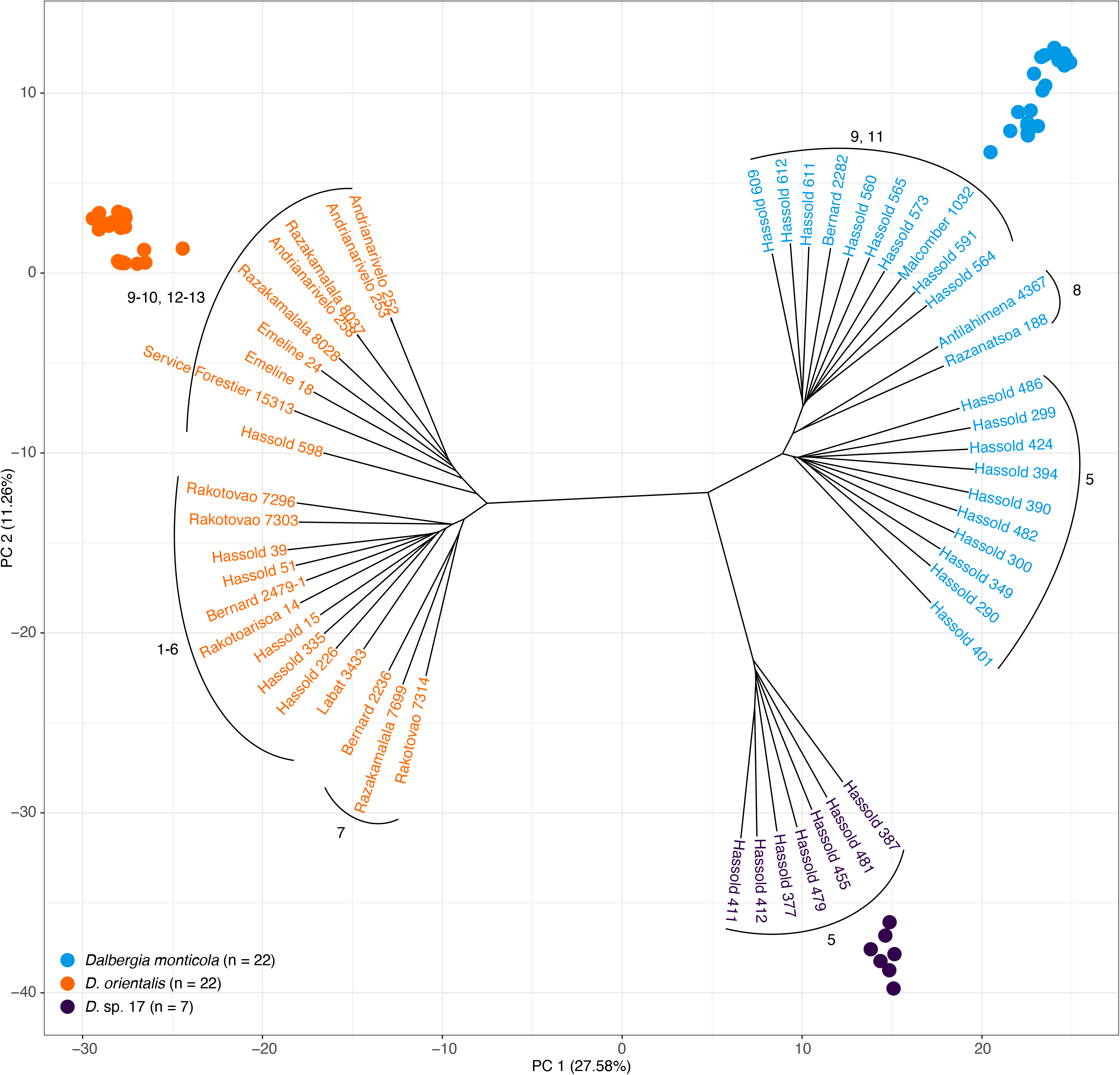
PCA and NJ tree of the population set (n = 51) inferred from 60,204 biallelic SNPs with no missing data. Numbers adjacent to tree branches denote sampling locations as shown in Figure S4.

## 4 DISCUSSION

Understanding the diversity and diversification of species and evolutionary lineages requires an integrative approach that links studies of micro-evolutionary processes to analyses of macro-evolutionary relationships (de La Harpe et al., 2017). Genetic data form a preferable source of information for investigations across broad evolutionary scales, as a large number of loci distributed across the nuclear genome can represent the spectrum of evolutionary rates at different scales of sample divergence. The present study introduces two overlapping sets of target capture probes for phylogenomic studies at micro-to macro-evolutionary timescales in rosewoods (Dalbergia2396 set) and across the legume family (Fabaceae1005 set), together with the flexible and modular bioinformatic pipeline CAPTUREAL, which streamlines the processing of sequencing reads for phylogenomic and population genomic analyses while visually informing users on the effect of critical parameter choices. We demonstrated the utility of individual assemblies per target region to produce alignments of hundreds of loci suitable for concatenation and multispecies coalescent approaches, which confirmed phylogenomic conflicts at the root of the legume family, and provided an unprecedented resolution of evolutionary relationships among lineages and species of the taxonomically complex genus *Dalbergia*. Remapping of sequencing reads further made it possible to identify thousands of informative sites amenable to population genomic analyses, which revealed the existence of a potentially new cryptic *Dalbergia* species. Together, these results illustrate that our newly developed probe sets are efficient tools for studies of species diversity and diversification in rosewoods (*Dalbergia* spp.) and more broadly in the economically important and highly diverse legume family.

### 4.1 Target capture probes

The target capture probes presented here are part of a growing collection of genomic resources for legume phylogenomics. Other probe sets for target capture in legumes have been developed, focusing on different groups within the family. Our probes can be compared to existing sets designed or validated at the level of legume species (Peng et al., 2017), genera (e.g., de Sousa et al., 2014; Nicholls et al., 2015; Shavvon et al., 2017), or subfamilies (Koenen et al., 2020a; Vatanparast et al., 2018) and across angiosperms (Johnson et al, 2019) to identify overlaps in target regions for legume phylogenomics. For example, it would be interesting to compare our probes with those of Vatanparast et al. (2018), who identified 423 target regions based on 30 transcriptomes, of which 27 were sampled from the NPAAA clade, one from Genistoids, and one each of the Caesalpinioideae and Cercidoideae subfamilies, limiting the probe set validation to 25 species of the NPAAA clade. Capture of additional, less conserved target regions across the legume family could be achieved by designing multiple probes for hybridization in the same target region in different legume groups, as applied for studies across angiosperms (Johnson et al., 2019). Such a probe design could profit from existing legume probe sets but should rely on a stringent selection of targets that accounts for paralogs (Vatanparast et al., 2018), which originated as a consequence of multiple whole-genome duplication events in legumes (Egan & Vatanparast, 2019; Koenen et al., 2021).

In this study, we enriched DNA libraries from three taxon sets spanning micro-evolutionary (populations) to macro-evolutionary (family) timescales, using a single set of 12,049 RNA probes targeting 6,555 genomic regions conserved across five Meso-Papilionoideae genomes and a *Dalbergia* transcriptome. We then identified 2,396 and 1,005 target regions with high capture specificity and sensitivity within the species-rich genus *Dalbergia* (Dalbergia2396 probe set) and more broadly across legumes (Fabaceae1005 probe set). We used our CAPTUREAL pipeline to refine phylogenomic and population genomic analyses using taxon-specific and longer reference sequences. This procedure has both benefits and drawbacks. An advantage is that different but overlapping probe sets amenable for efficient target capture in different focal groups can be identified, and that a single enriched DNA library can be included in multiple data sets spanning different evolutionary timescales. On the other hand, bioinformatic analyses took longer due to the iterative refinement, and only a portion of captured sequence data was ultimately used for phylogenomic or population genomic analyses in each focal group (see Tables S1 – S3). Higher costs per used sequence could be compensated by enriching DNA of up to six individuals in a single hybridization reaction, a strategy that has been used successfully in other studies (e.g., de La Harpe et al., 2018; Yardeni et al., 2021).

### 4.2 CAPTUREAL bioinformatics pipeline

The CAPTUREAL pipeline starts with the mapping of quality-trimmed reads to conserved target regions identified during probe design, followed by assembly on a per-region basis, orthology assessment, and filtering for target regions with high capture sensitivity and specificity for downstream analyses. This approach differs from the PHYLUCE pipeline (Faircloth, 2016), where quality-trimmed reads are first assembled, and then matched to target regions. CAPTUREAL simplifies the assembly of reads specific to each locus, circumventing the challenging task of *de novo* assembly of contigs from the large pool of sequencing reads representative of thousands of loci (reviewed by Chaisson et al., 2015). Likewise, alignments are conducted in clearly defined target regions in which overlap among individual contigs is higher. However, assembly per region is more time-consuming and requires reference sequences for the initial mapping step. This might introduce a reference bias when divergent sequences are not mapped (Lunter & Goodson, 2011). We addressed this problem by generating consensus sequences that are representative of a given taxon set and by limiting analyses to target regions that can be efficiently recovered in all groups of that taxon set. These set-specific reference sequences can then be used to iteratively refine mapping, assembly, and target region filtering for any set of taxa. Our approach is conceptually similar to the HYBPIPER pipeline (Johnson et al., 2016), which also employs a mapping-assembly strategy, and uses depth of coverage and percent identity to the target region to choose between multiple contigs, before it identifies intron/exon boundaries using target peptide sequences and extracts coding sequences for alignment. While the HYBPIPER pipeline is designed specifically for the HYB-SEQ approach (Johnson et al., 2016), in which exons are the primary targets and flanking non-coding regions are used as supplementary data for analyses at shallow evolutionary scales (Weitemier et al., 2014), CAPTUREAI is more general in scope and neither requires nor leverages knowledge about intron/exon boundaries in the targeted regions. It is therefore suitable for application in systems lacking high-quality annotated reference genomes or transcriptomes. The main strengths of this pipeline are its modularity, which allows for an iterative refinement of read mapping, assembly and alignment, its flexibility given by user-defined parameters, the merging of alignments representing physically overlapping target regions, and the visualization of key summary statistics and alignments along the workflow to inform the user on critical analysis parameters.

### 4.3 Macro- and micro-evolutionary patters in *Dalbergia*

*Dalbergia* species endemic to Madagascar were recovered as two large, well-supported and fully resolved clades, each exclusively comprising Malagasy species. These two clades were previously identified on the basis of three chloroplast markers, but phylogenetic relationships within clades were not resolved, which exposed traditional DNA barcoding as insufficient for genetic discrimination between closely related *Dalbergia* species (Hassold et al., 2016; see Tables S2 and S3). Supergroup I and II are morphologically divergent and largely correspond to Group 1 and 2 reported by Bosser & Rabevohitra (2002). Supergroup I is characterized by a glabrous gynoecium with a long and slender style and relatively large flowers, while Supergroup II is characterized by a pubescent gynoecium with a short and squat style and relatively small flowers. The two supergroups are both more closely related to non-Malagasy taxa than to each other, suggesting at least two independent colonizations of Madagascar followed by species diversification. The only sampled Malagasy species not belonging to either of the two supergroups is *D. bracteolata*, which occurs on Madagascar as well as in mainland East Africa. A further species, which is endemic to Madagascar and morphologically divergent from Supergroups I and II (*D. xerophila* Bosser & R. Rabev.) was not included in this study.

Within Supergroup I, two well-supported subclades were resolved, which differ in their inflorescence structure. Within *Dalbergia chapelieri* s.l., a widely distributed species complex with paniculate inflorescences, northeastern and southeastern populations can be distinguished using the present data as well as chloroplast variation (Hassold et al., 2016). The other subclade within Supergroup I contains species from eastern Madagascar with mostly racemose inflorescences, including a potentially new species, *Dalbergia* sp. 24. Collections belonging to this entity were previously believed to be conspecific with *D. maritima* var. *pubescens* (see Hassold et al., 2016) but show geographic (i.e., north-east vs. central-east), morphological (i.e., more numerous leaflets that are smaller, more oblong and less coriaceous) and genetic (Figure 2, Figure S3) differences compared to the type material (*Service Forestier 32824*). The type (collected in 1985) showed a slightly longer terminal branch compared to other samples in the concatenation tree (Figure S3) but clearly grouped with two recently collected conspecific samples from central-east Madagascar. The same subclade also contains material of two highly valued rosewood species, *D. occulta* and *D. normandii* (note that in Hassold et al. (2016), sterile material of *D. normandii* was erroneously identified as *D. madagascariensis*).

Supergroup II includes two clades distributed in the humid and sub-humid east and northwest of Madagascar, and a large third clade centered in the drier west and north of the island. Morphological synapomorphies characterizing these clades require further genetic and morphological analyses. The geographic separation in major eco-geographic regions of Madagascar suggests that climate regimes may have played a significant role in shaping the evolution of these groups, which thus constitute promising model systems to study processes of ecological divergence, along the same lines as studies that have investigated elements of the Malagasy fauna (Vences et al., 2009).

Our results revealed relationships among Supergroups I and II and non-Malagasy taxa that are incompatible with the plastid phylogeny of Hassold et al. (2016), in particular with regard to *Dalbergia melanoxylon* (Africa), *D. ecastaphyllum* (America and Africa), and *D.* cf. *oliveri* (Asia). Incongruence between nuclear and plastid phylogenies is common at various evolutionary timescales in many plant groups (e.g., Lee et al., 2021; Pelser et al., 2010), and while the multispecies approach applied in this study is expected to return a phylogeny that reflects nuclear evolution accounting for incomplete lineage sorting, conflicts in gene tree topologies due to hybridization and chloroplast capture can further underlie the observed differences.

Our target capture approach demonstrated great potential to facilitate the resolution of several taxonomic conundrums within the genus, which likely resulted from few observable and diagnostic morphological characters, insufficient collection effort, and the difficulty of distinguishing between heritable and plastic trait variation within and among *Dalbergia* species (Lachenaud, 2016). The integration of highly informative museum specimens, including a nomenclatural type collected in 1985, enabled the accurate identification of recently collected but often sterile specimens, and was crucial in detecting misidentifications or potential taxonomic inadequacies (Buerki & Baker, 2016), as shown for *D. maritima* var. *pubescens* or *D. monticola*.

Population genomic analyses of 51 individuals readily separated the two closely related species *Dalbergia monticola* and *D. orientalis*, as well as a sympatric and syntopic but genetically differentiated entity, which could previously not be differentiated from the other two species based on three chloroplast markers (Hassold et al., 2016). The lack of admixture between *D. monticola* and this third cluster, the similarity in leaf characters, and the absence of known morphologically similar species occurring in the region, prompts us to hypothesize the latter to reflect a separate, yet undescribed cryptic species. Both *D. monticola* and *D. orientalis* are distributed from northeastern to south-eastern Madagascar, co-occur in various localities, but differ in their predominant altitudinal distribution (Madagascar Catalogue, 2021). Population structure within both species was uncovered using our target capture approach and appears to be sufficient to distinguish specimens from the northeast (locations 1 to 6), central-east (locations 7 and 8), and southeast of the island (locations 9 to 13). These results indicate that genetic species identification and provenancing, at least to this broad geographic scale, may be feasible, which would have important implications for forensic timber identification and for tracing geographic hotspots of the illegal trade in these valuable timber species (UNODC, 2016a).

### 4.4 Phylogenetic analyses across legumes

At the family level, 1,005 merged regions of the 6,555 targeted regions passed our stringent sensitivity and specificity filters, suggesting that many target regions were not efficiently captured across taxa. However, phylogenetic analysis of 986 nuclear target regions recovered multiple known clades within monophyletic subfamilies with strong bootstrap and quartet support, providing excellent resolution comparable to that obtained in the recent nuclear phylogenomic analysis of transcriptome and genome-wide data across legumes (Koenen et al., 2020b). As in that study, we found high support for Cercidoideae and Detarioideae as sister taxa, a relationship that was never inferred in analyses based on chloroplast genes (LPWG, 2017) or plastomes (Koenen et al., 2020b; Zhang et al., 2020). As shown in both studies, the other relationships among subfamilies are difficult to resolve. Our most supported subfamily topology (38.4% quartet support, Figure S2A) recovered the Papilionoideae as sister to a clade comprised of Caesalpinioideae, Dialioideae, and the Cercidoideae/Detarioideae clade, while Koenen et al. (2020b) demonstrated a successive divergence of the Cercidoideae/Detarioideae clade, Dialioideae, Caesalpinioideae and Papilionoideae in all nuclear analyses. This alternative topology received almost equivalent overall quartet support (38.36%) in our analyses (Figure S2C), as did a third hypothesis in which Caesalpinioideae and Papilionoideae are sister to Dialioideae and the Cercidoideae/Detarioideae clade (Figure S2B). These nearly equally supported subfamily topologies can be explained by short deep internodes associated with conflicting bipartitions and are consistent with the idea of a nearly simultaneous evolutionary origin of all six legume subfamilies, causing incomplete lineage sorting (Koenen et al., 2020b). Taxon sampling may additionally contribute to the contentious deep-branching relationships. The monotypic Duparquetioideae subfamily could not be analyzed, and a portion of gene trees may suffer from long branch attraction between *Polygala* and Papilionoideae, which both exhibit markedly higher substitution rates compared to the other legume subfamilies (Koenen et al., 2020b). Additional outgroup taxa such as members of the Quillajaceae family could alleviate this problem, and permit a more accurate inference of subfamily relationships.

Substantial gene tree incongruence was also found with respect to the relationships among the three large clades within Meso-Papilionoidae. The sister relationship between Dalbergioids s.l. and Genistoids s.l. received only slightly higher quartet support than the two alternative hypotheses, which is consistent with previous results (Koenen et al., 2020b). Similarly, conflicting topologies affected most branches within Genisteae. By contrast, our analyses confirm that the genus *Aeschynomene* sensu RUDD (1955), which consisted of the former *A.* sect. *Aeschynomene* and *A.* sect. *Ochopodium* Vogel, is non-monophyletic (Ribeiro et al., 2007). The recently re-established *Ctenodon* (= *A.* sect. *Ochopodium*, Cardoso et al., 2020) is sister to *Machaerium*, and these two genera form the sister group to *Dalbergia*.

### 4.5 Conclusions and perspectives

The resources developed here for Fabaceae and in particular the genus *Dalbergia* bridge micro- and macro-evolutionary timescales and will hopefully facilitate community-driven efforts to advance legume genomics. Comprehensive sampling and sequencing by target capture of *Dalbergia* across its distribution range, and in particular from the hotspot of diversity in Madagascar, can yield valuable insights into the origin and diversification of the genus, thereby informing conservation policies and the taxonomic revision of Malagasy *Dalbergia*. The obtained sequence data will further serve to build a reference library for molecular identification of CITES-listed *Dalbergia* species, which would make a significant contribution toward the conservation of the valuable and endangered rosewoods.

## ACKNOWLEDGEMENTS

We thank Sonja Hassold and the Missouri Botanical Garden (MBG) Madagascar team for organising and conducting field work, the Botanic Garden of the University of Zurich for the opportunity to sample legume plants from their living collection, and to MBG in St. Louis (USA) and the Paris herbarium (P) for access to their DNA bank and voucher specimens, respectively. We also thank Peter Phillipson and Nicholas Wilding for discussing sample determinations, *Dalbergia* taxonomy and Malagasy plant diversity. We are grateful to Claudia Michel for laboratory work and the Genetic Diversity Centre Zurich (GDC) for helpful support, in particular to Silvia Kobel for sequencing and Niklaus Zemp for bioinformatics. Finally, we thank Erik Koenen, Colin Hughes, Pete Lowry, Martin Fischer and Nicholas Wilding for their valuable inputs and comments on the manuscript.

## AUTHOR CONTRIBUTIONS

SC and AW designed the study and collected samples. SZ assembled the draft *Dalbergia* transcriptome and designed target capture probes. SC and SZ analyzed data and wrote CAPTUREAL. SC, SF and AW wrote the manuscript with contributions from SZ.

## DATA AVAILABILITY STATEMENT

Raw target capture sequencing reads generated for this study are deposited in the European Nucleotide Archive (ENA) at EMBL-EBI under accession number PRJEB41848 (https://www.ebi.ac.uk/ena/browser/view/PRJEB41848). Transcriptome sequencing reads as well as the draft *Dalbergia* transcriptome, sequences representing the initial 12,049 RNA probes and 6,555 target regions, the Fabaceae1005 and Dalbergia2396 probe sets, longer and taxon-specific reference sequences used for mapping, final alignments for the subfamily and species sets (all in FASTA format), and SNP data from the population set (VCF format) are available on Dryad (https://doi.org/10.5061/dryad.73n5tb2z7). The bioinformatics pipeline CAPTUREAL is available and further documented on Github (https://github.com/scrameri/CaptureAl). Because *Dalbergia* species are under threat from illegal exploitation, we have systematically refrained from making detailed distribution maps and precise geo-coordinates publicly available. Specimen records for collections from Madagascar are provided in the Catalogue of the Plants of Madagascar (Madagascar Catalogue, 2021), but with restricted public access to precise geo-coordinates (delivered on demand to bona fide researchers).

