## Supplemental Information for "A target capture approach for phylogenomic analyses at multiple evolutionary timescales in rosewoods (*Dalbergia* spp.) and the legume family (Fabaceae)"

#### Table of Contents:


| | |
|---|---|
| **Supplementary Figures S1 – S9** | Pages 2 – 10 |
| **Supplementary Tables S1 – S7** | Page 11 |
| **Supplementary Methods** | Pages 12 – 22 |
| **Supplementary References** | Page 23 |

### Supplementary Figures

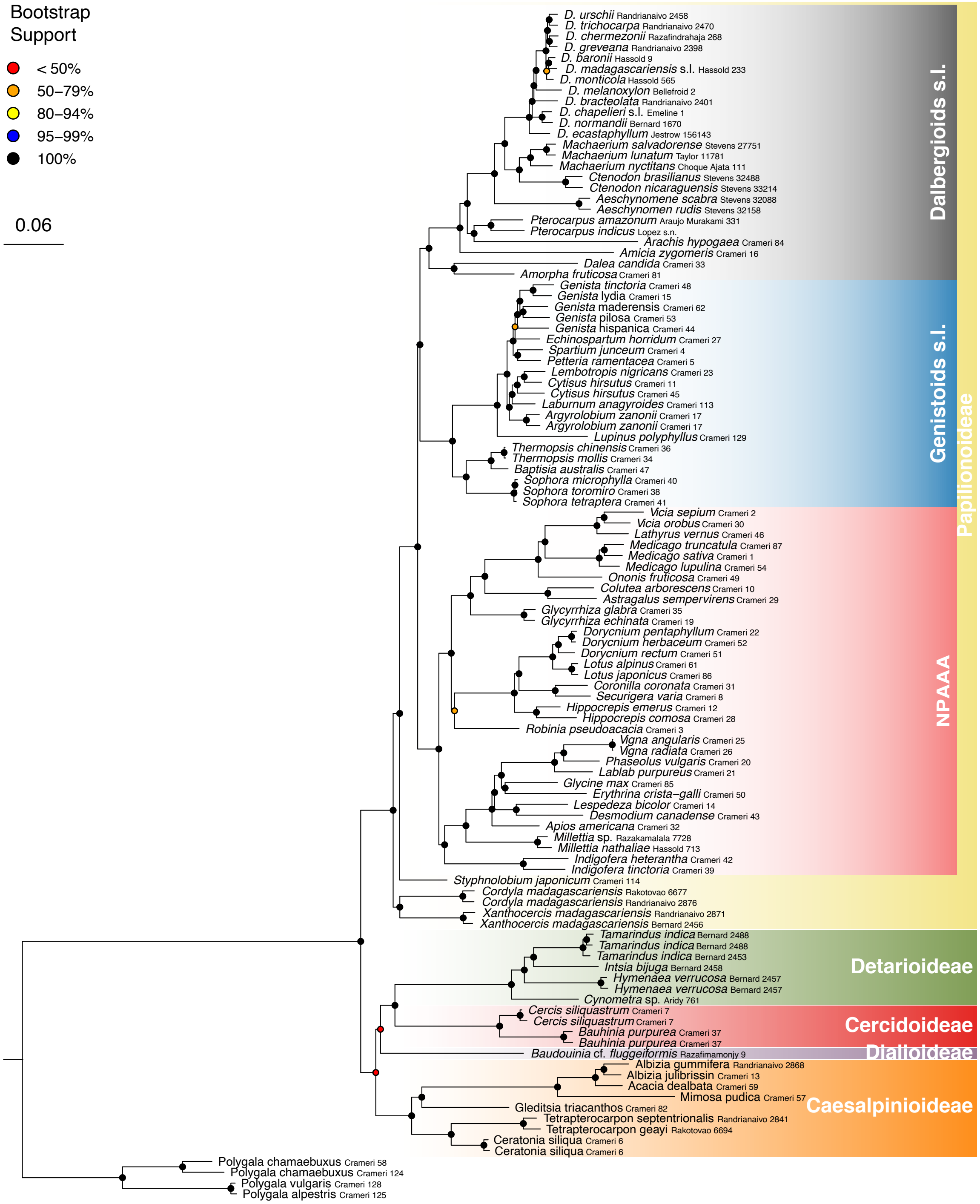


FIGURE S1 Maximum likelihood tree of the subfamily set (n = 110) inferred using RAxML on the alignment supermatrix (concatenation method). The supermatrix had 1,196,506 alignment sites, 1,028,714 distinct alignment patterns (unique columns), and 21.73% missing data.

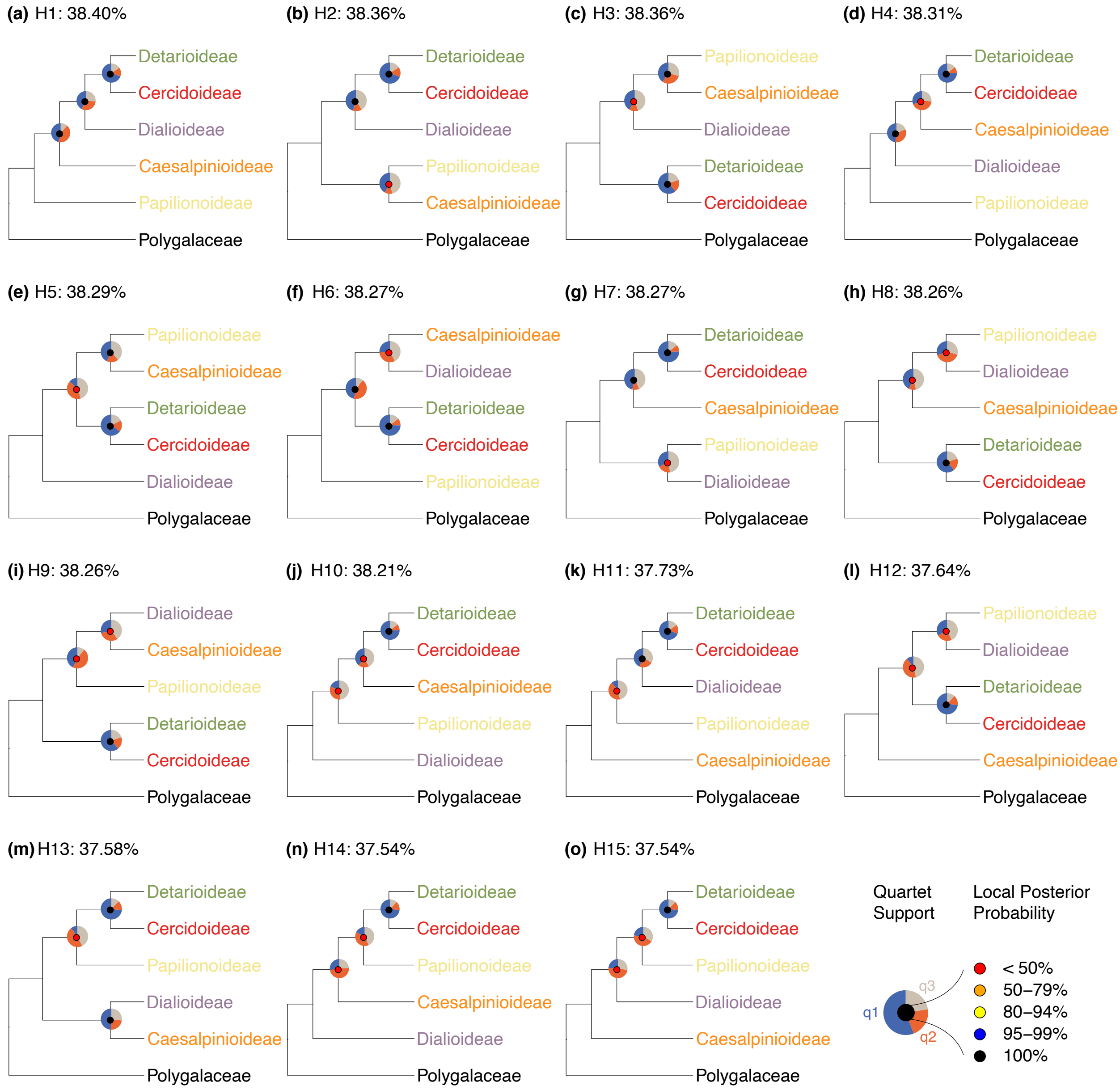


FIGURE S2 Quartet support for fifteen hypotheses (H1 to H15) on relationships among Fabaceae subfamilies. Panels (a) to (o) each represent one hypothesis given by one topology. Panels are ranked by decreasing normalized quartet score computed by ASTRAL-III, i.e., the percentage of input gene tree quartet trees that are consistent with a topology. Pie charts on each node denote the fraction of gene trees that are consistent with the shown topology (q1; blue), and with the first (q2; orange) and second (q3; gray) alternative topologies. Local posterior probabilities are shown as color-coded circles on each node (see legend).

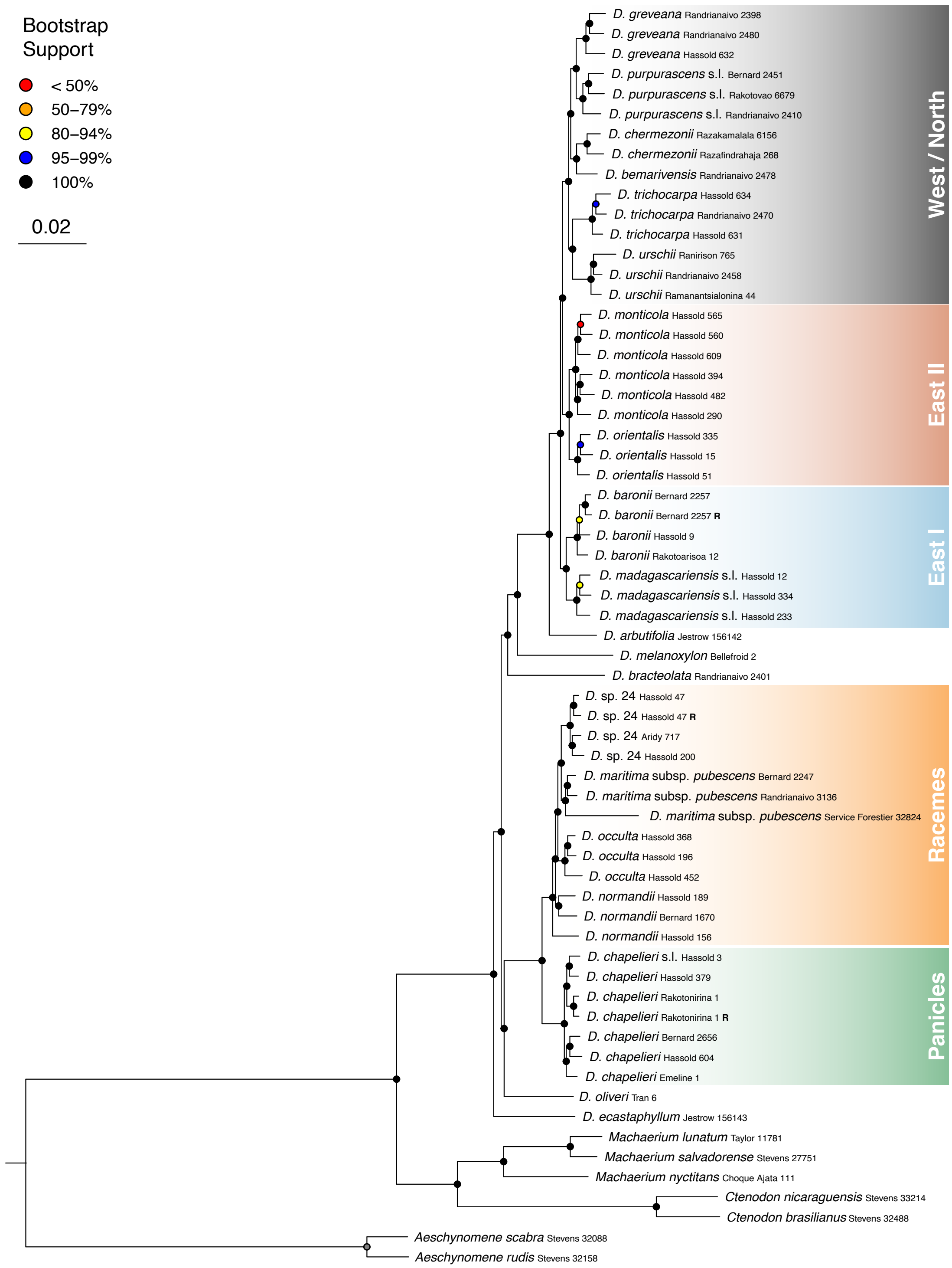


FIGURE S3 Maximum likelihood tree of the species set (n = 63) inferred using RAxML on the alignment supermatrix (concatenation method). The supermatrix had 2,589,455 alignment sites, 1,270,746 distinct alignment patterns (unique columns), and 17.18% missing data.

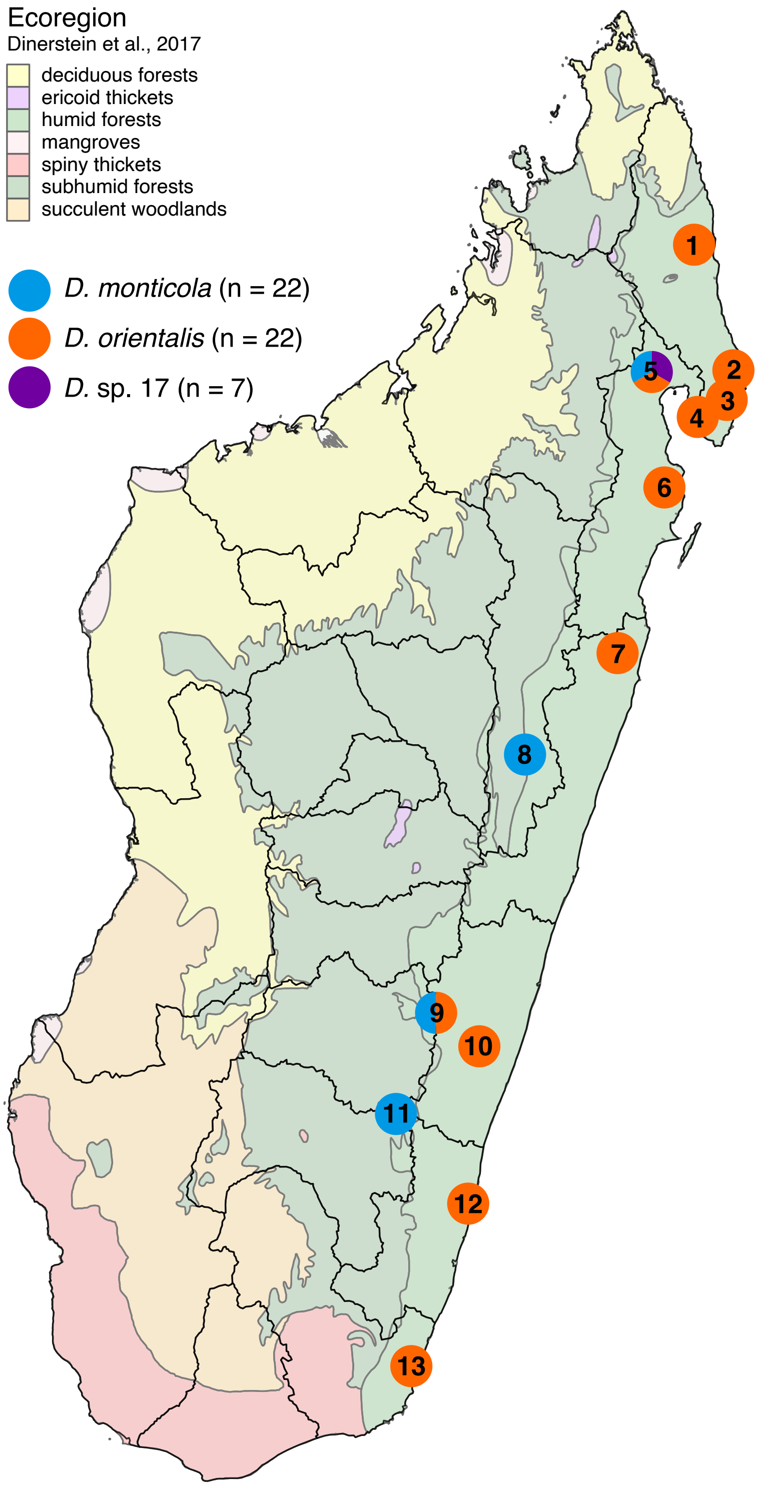


FIGURE S4 Sampling locations of species set specimens (n = 51). Pie charts denote the proportion of samples of each species collected at each location. See numbers on pie charts and Table S3 for details on specimens collected at each location. The map was created using TMAP version 3.0 (Tennekes, 2018).

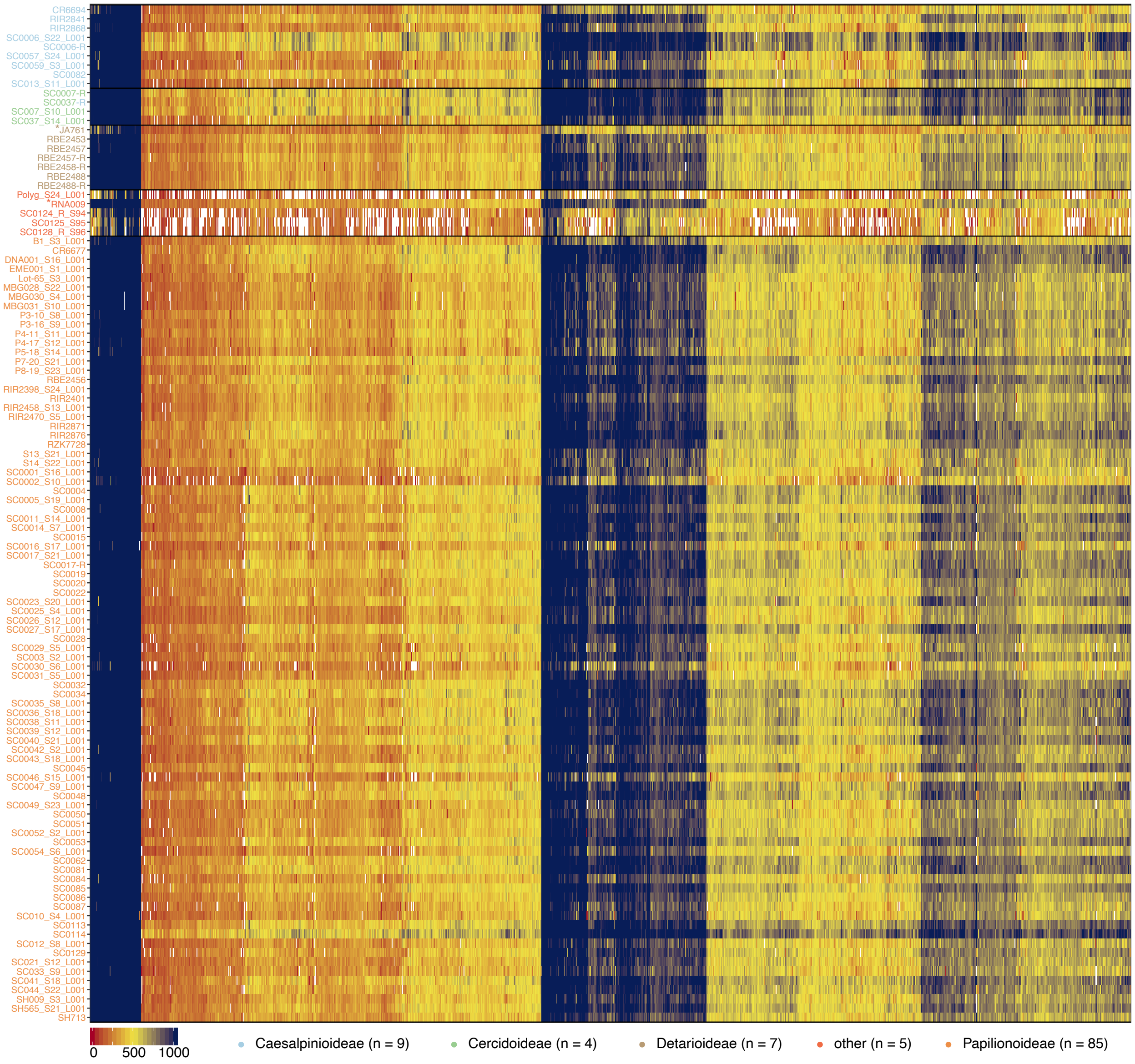


FIGURE S5 Heatmap of covered length in base pairs in 1,917 target regions (x axis) retained after step 2 of the second iteration in the subfamily set specimens (y axis, n = 110). Specimens are sorted according to the color-coded taxon groups used for target region filtering. Target regions are sorted according to hierarchical clusters. Values above 1,026 base pairs were capped for better readability. Sample identifiers are as in Table S1. Two samples marked with asterisks were extracted from herbarium vouchers.

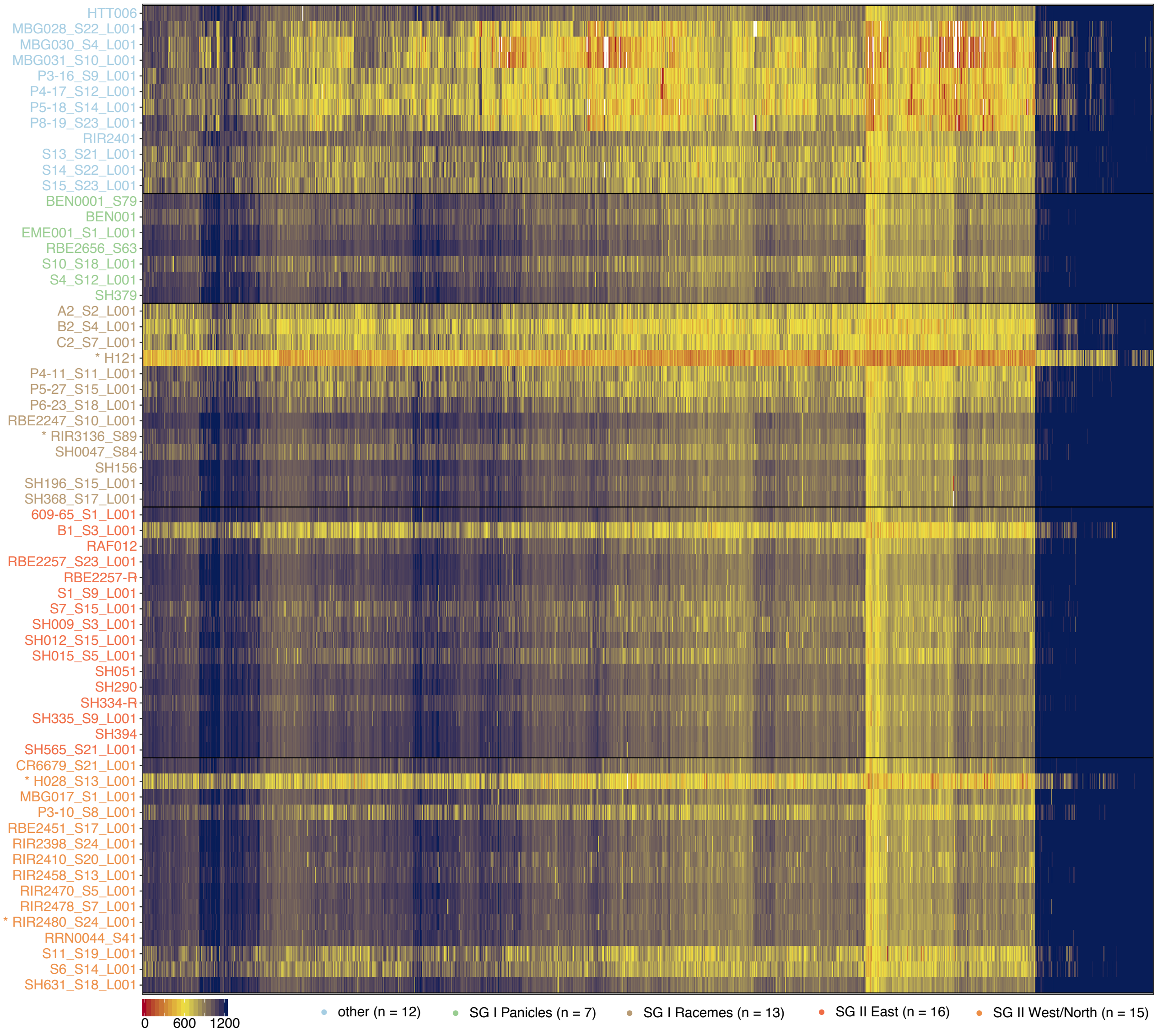


FIGURE S6 Heatmap of covered length in base pairs in 3,418 target regions (x axis) retained after step 2 of the second iteration in the species set specimens (y axis, n = 63). Specimens are sorted according to the color-coded taxon groups used for target region filtering. Target regions are sorted according to hierarchical clusters. Values above 1,221 base pairs were capped for better readability. Sample identifiers are as in Table S2. Four samples marked with asterisks were extracted from herbarium vouchers.

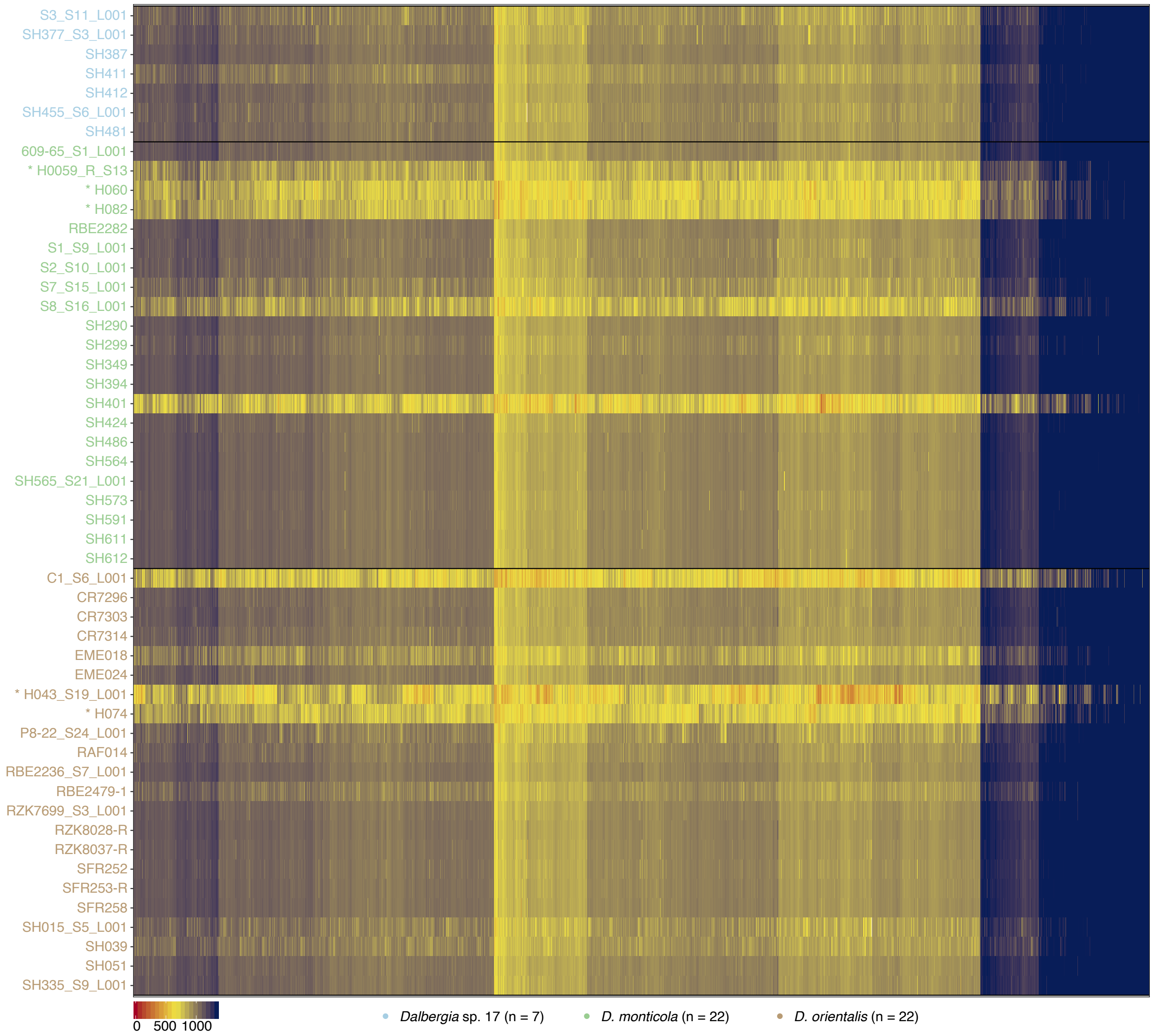


FIGURE S7 Heatmap of covered length in base pairs in 2,396 target regions (x axis) used as mapping targets in the species set specimens (y axis, n = 51). Specimens are sorted and color-coded for species. Values above 1,339 base pairs were capped for better readability. Target regions are sorted according to hierarchical clusters. Sample identifiers are as in Table S3. Five samples marked with asterisks were extracted from herbarium vouchers.

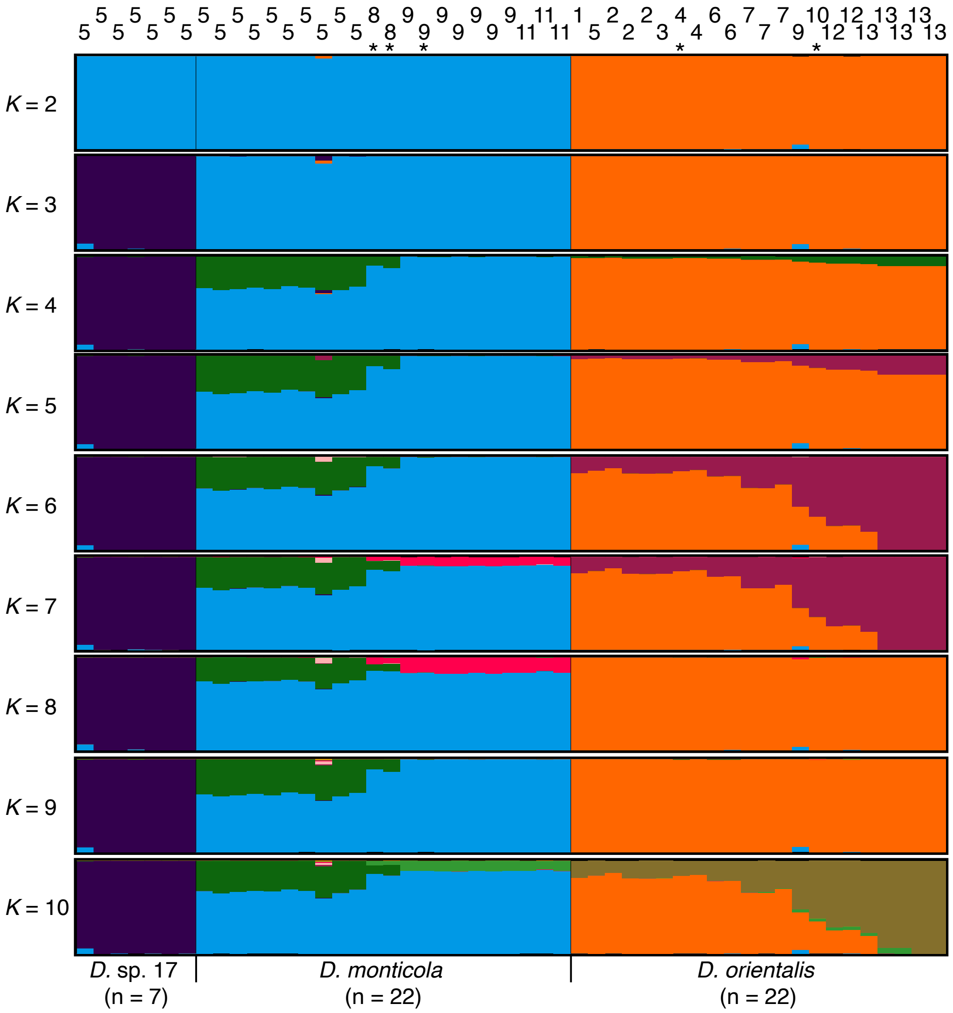


FIGURE S8 Structure results for up to ten assumed clusters *K* in 51 individuals and 7,156 single nucleotide polymorphisms (SNPs). Individuals are sorted by species and then by increasing degrees south latitude. Numbers at the top indicate broad sampling locations as in Figure S4 and Table S3. The major clusters averaged across ten replicate runs using CLUMPAK (Kopelman et al., 2015) are shown. The first split at *K* = 2 separated *Dalbergia orientalis* from *D. monticola* and *D.* sp. 17, and the second split at *K* = 3 further separated *D. monticola* from *D.* sp. 17. Admixture proportions were less clear-cut for higher values of *K*, but indicated isolation by distance at a broad geographical scale, dividing specimens from northeast (locations 1 to 6), central-east (locations 7 and 8) and southeast Madagascar (locations 9 to 13) in both *D. monticola* and *D. orientalis*. Samples marked with an asterisk (*) denote were extracted from herbarium vouchers.

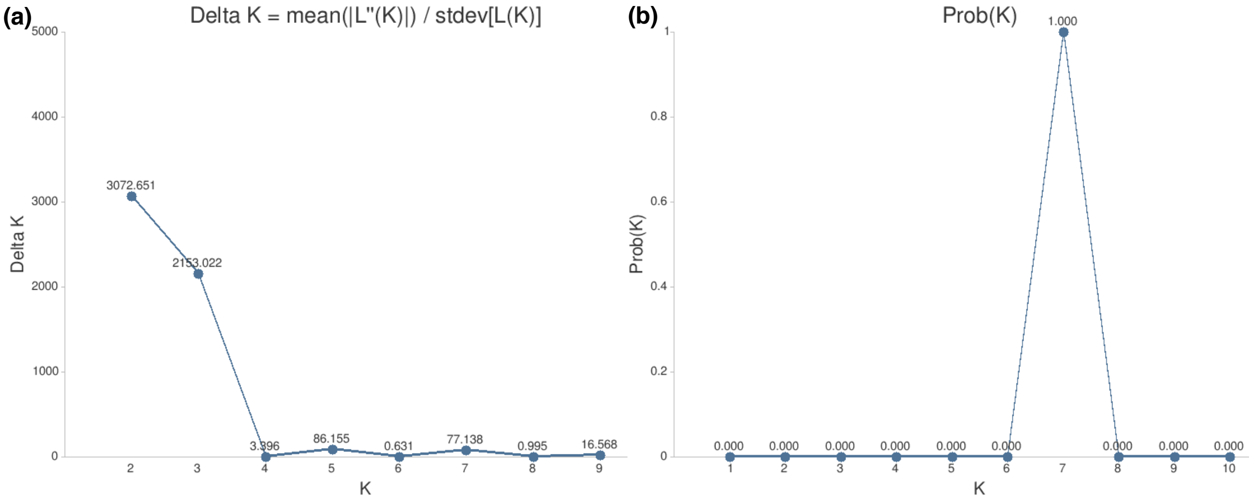


FIGURE S9 Structure probability at different values of *K*. (a) Delta *K* statistic. (b) Probability by *K.* Graphs were produced using CLUMPAK (Kopelman et al., 2015).

#### Supplementary Tables

Supplementary Tables S1 – S5 are available as a single Excel file with five sheets.

- TABLE S1 Collection details (turquoise columns), sequencing statistics (gray columns), mapping statistics (orange columns) and assembly statistics (blue columns) of the subfamily set (n = 110).

- TABLE S2: Collection details (turquoise columns), sequencing statistics (gray columns), mapping statistics (orange columns) and assembly statistics (blue columns) of the species set (n = 63).

- TABLE S3: Collection details (turquoise columns), sequencing statistics (gray columns) and mapping statistics (orange columns) of the population set (n = 51).

- TABLE S4 Characterization of genomic regions targeted by the Fabaceae1005 probe set.

- TABLE S5 Characterization of genomic regions targeted by the Dalbergia2396 probe set.

Supplementary Tables S6 – S7 are presented in Supplementary Methods (pp. 12 – 23).

### Supplementary Methods

#### Assembly of a draft *Dalbergia* transcriptome

For transcriptome assembly, we collected fresh leaf material of *Dalbergia madagascariensis* Vatke s.l. from a young plant (progeny of collection Jean-Luc Mora 26 from the Masoala peninsula) cultivated in a greenhouse at ETH research station Lindau-Eschikon, Switzerland. Young leaves and leaf buds were snap-frozen in liquid nitrogen and stored until used. Frozen leaves were later ground with a BeadRuptor (Omni International) and total RNA was isolated using the RNeasy® Plant Mini Kit (Qiagen). We quantified total RNA using the RNA BR (broad range) assay kit (Life Technologies) for QUBIT™ 2.0 fluorometer (Promega), checked RNA integrity on a 1.5% agarose gel, and verified RNA quality on an Agilent 2100 Bioanalyzer. Total RNA was further treated with DNase I prior to library preparation with the Illumina® TruSeq RNA kit (Illumina) at the Functional Genomics Centre, Zurich, Switzerland (FGCZ). Paired-end sequencing was performed on an Illumina HiSeq™ 2000. We obtained 63 million paired-end reads, corresponding to a total of 6.2 gigabases (Gb). The raw reads were cleaned by removing adapter sequences with CUTADAPT (Martin, 2011), followed by filtering and trimming of low-quality reads and bases *with condetri.pl* (Smeds & Künstner, 2011). We performed a de novo assembly of the transcriptome using TRINITY release 2012-01-25 (Grabherr et al., 2011), resulting in 146,484 scaffolds that were between 201 and 17,129 bp long, with a mean length of 815 bp. Raw sequence data and the draft *Dalbergia* transcriptome are deposited on Dryad (https://doi.org/10.5061/dryad.73n5tb2z7).

#### Design of target capture probes and reference sequences

We selected high-quality genome assemblies of five legume species available in public databases for divergent reference capture (Jones & Good, 2016): *Cajanus cajan* (L.) Millsp. v1.0 (https://www.ncbi.nlm.nih.gov/assembly/GCF_000340665.1), *Glycine max* (L.) Merr. v1.0 (http://phytozome.jgi.doe.gov), *Lotus japonicus* L. v2.5 (https://lotus.au.dk/data/download), *Medicago truncatula* Gaertn. v3.5 (https://www.ncbi.nlm.nih.gov/assembly/GCF_000219495.1), and *Phaseolus vulgaris* L. v2.1 (DOE-JGI and USDA-NIFA, http://phytozome.jgi.doe.gov). We pairwise aligned the draft *Dalbergia* transcriptome and the five genome sequences using LASTZ (Harris, 2007) to extract 60-bp long sequences that were fully conserved across species. After removing direct and reverse-complement duplicates from 169,484 sequences, we retained 74,604 sequences. We then mapped the sequences back to each genome/transcriptome using BLAST (Altschul et al., 1990) and kept 10,867 sequences that returned only one ungapped hit after filtering for a minimum of 90% overlap and 95% identity in all genomes. We processed the resulting sequencing conditional to their length by 1) dividing sequences longer than 150 bp into 100 bp probes with a 50 bp overlap; 2) extracting a central probe of 100 bp for sequences measuring 100 to 150 bp in length; 3) extending shorter sequences to 100 bp based on the *C. cajan* genome; or 4) discarding sequences that could not be extended, e.g. because they were at the edge of an assembly scaffold. After removal of duplicates and reverse complement duplicates, we removed probes with low complexity and interspersed repeats using REPEATMASKER (Smit et al., 2013) and further discarded those with a GC-content higher than 70%. Following a further BLAST run and filtering with relaxed settings (one hit, overlap >= 85%, identity >= 92.5%), we retained a final set of 12,049 partially overlapping probes from 7,201 conserved regions that we ordered for synthesis as myBaits Custom Target Capture Kits (Arbor Biosciences; https://arborbiosci.com).

To generate a reference sequence for the bioinformatics analyses, we combined sequences of regions that were within 100 bp of one another by filling the gaps based on the *Cajanus cajan* assembly. This resulted in reference sequences for 6,555 regions (target regions hereafter), with a mean length of 188 bp (100 to 1,424 bp), a total length of 1,233,157 bp, and a mean physical distance of 52,802 bp (100 to 1,306,862 bp) between sequences of the same genomic scaffold. The target regions covered all eleven linkage groups of the *C. cajan* assembly (3,931 target regions, 104 to 761 target regions per linkage group) while 2,624 target regions belonged to unanchored genomic scaffolds (1 to 48 target regions per scaffold).

**Taxon samples for target capture probes validation**

The eight species of Caesalpinioideae included four species of the mimosoid clade (LPWG, 2017). The 85 Papilionoideae samples were represented by one species of the Cladrastis clade (Wojciechowski, 2013), two species of the Angylocalyx-Dipterygeae-Amburana (ADA) clade (Cardoso et al., 2012) and 78 species of Meso-Papilionoideae (Wojciechowski, 2013). Within the Meso-Papilionoideae, we included representatives of three species-rich clades: 19 species of Genistoids s.l. (Wojciechowski et al., 2004; Cardoso et al., 2012), 25 species of Dalbergioids s.l. (Wojciechowski et al., 2004), and 34 species of the non-protein-amino-acid-accumulating (NPAAA) clade (Wojciechowski et al., 2004; Cardoso et al., 2012).

Samples were obtained from various sources. We re-analyzed 46 DNA samples from a study on *Dalbergia* chloroplast variation (Hassold et al., 2016), analyzed 39 *Dalbergia* samples collected for this study in Madagascar, and 48 samples collected with permission from the Zurich Botanical Garden (Switzerland). We also obtained twelve samples through the Missouri Botanical Garden DNA bank (St. Louis, MO, USA), three herbarium samples from the Conservatoire et Jardin botaniques Genève (CJBG), three herbarium samples from the Muséum National d'Histoire Naturelle (MNHN) Paris, and one sample from the USDA-ARS Tropical Agriculture Research Station (Mayaguez, Puerto Rico). Ten samples were collected in the field in Switzerland by S. Crameri, A. Widmer and M. Baltisberger. Five samples were grown from commercially available seeds at the ETH research station Lindau (Switzerland) and two samples were purchased from a commercial source.

**Read quality-trimming and quality-filtering**

Raw paired-end reads with the indicated file extensions (`-x` option) located in `$rawreads` (`-r` option) were quality-trimmed and quality-filtered using trimmomatic version 0.32 (Bolger et al., 2014). Specifically, we used ILLUMINACLIP with an adapter sequence file containing NEBNext, TruSeq and Illumina adaptor sequences (-a option), a seed mismatch of 2, a palindrome clip threshold of 20, a simple clip threshold of 10, a minimum adapter length of 10, while keeping both reads. Leading and trailing bases of each read were removed if the quality was below 5. Sliding window trimming was performed using a window size of 4 and a required average quality of 15. Quality-trimmed reads shorter than 50 bases were removed. The following script executed trimmomatic as specified above, for 20 samples in parallel:

```
trim.fastq.sh -s $s -a illumina.truseq.indexing.adaptors -r $rawreads -x
'_R1.fastq.gz,_R2.fastq.gz' -t 20
```

The quality of raw and trimmed reads was assessed with FastQC version 0.11.5 (https://www.bioinformatics.babraham.ac.uk/projects/fastqc).

**Executed command-line scripts and parameter choices for steps 1–7 and iterations 1–2**

The following tables and sections track the executed pipeline scripts and chosen parameters at each analysis step. For clarity, executed scripts are only listed once despite having been executed for several taxon sets and iterations. Parameter choices that varied between taxon sets or iterations are denoted as `$X` and are specified in Tables S6 and S7, respectively. Pipeline scripts are further documented on the README.md and wiki pages on the accompanying GitHub page (https://github.com/scrameri/CaptureAl).

TABLE S6 CaptureAl parameter choices for the subfamily set. Two separate columns denote parameters chosen during Steps 1–7 of the first and second iteration of mapping, assembly and target region alignment. Other pipeline parameters are case-specific number of threads for parallel computation (`-t`), names of folders containing output of previous analysis steps, or parameters left at their default values. nb. = number; frac. = fraction; avg. = average; aln. = alignment; min. = minimum; max. = maximum.

| Step(s) | Var | Parameter | Iteration 1 | Iteration 2 |
|---|---|---|---|---|
| 1+2+3+6 | $s | sample file | samples.fabaceae.12.txt | samples.fabaceae.txt |
| 1+3+4 | $1 | reference sequences | Cajanus_cajan_6555reg.fasta | Fabaceae_iter1_2468reg.fasta |
| 1 | $T | bwa-mem alignment score | 10 | 10 |
| 1+2 | $Q | min. mapping quality | 10 | 10 |
| 1 | $e | trimmed read file extension | .trim1.fastq.gz,.trim2.fastq.gz | .trim1.fastq.gz,.trim2.fastq.gz |
| 1+4+5+7 | $2 | sample / taxon group file | mapfile.fabaceae.12.txt | mapfile.fabaceae.txt |
| 1 | $3 | min. frac. regions | 0.2 | 0.2 |
| 1 | $4 | min. frac. taxa | 0.3 | 0.7 |
| 1+3 | $5 | min. aln. length | 1 | 1 |
| 1 | $6 | min. avg. coverage | 6 | 8 |
| 1 | $7 | max. avg. coverage | 1000 | 1000 |
| 1 | $8 | min. aln. fraction | 0 | 0 |
| 1 | $9 | min. frac. conforms | 0.3 | 0.7 |
| 3 | $10 | min. normalized score | 2 | 1 |
| 4+5 | $11 | min. frac. regions | 0.2 | 0.2 |
| 4+5 | $12 | min. frac. taxa | 0.5 | 0.75 |
| 4+5 | $13 | max. nb. contigs | 2 | 2 |
| 4+5 | $14 | min. normalized score | 2 | 2 |
| 4+5 | $15 | min. aln. length | 80 | 80 |
| 4 | $16 | min. aln. fraction | 0 | 0 |
| 4 | $17 | min. score | 1 | 1 |
| 4 | $18 | min. contig length | 1 | 1 |
| 4+5 | $19 | min. frac. conforms | 0.5 | 0.5 |

| | | | | |
|---|---|---|---|---|
| 6 | $20 | min. merging score | 0.85 | 0.85 |
| - | $21 | max. gap ratio | - | 0.35 |
| - | $22 | max. nucleotide diversity | - | 0.35 |

TABLE S7 CaptureAl parameter choices for the species set. Two separate columns denote parameters chosen during Steps 1–7 of the first and second iteration of mapping, assembly and target region alignment. Other pipeline parameters are case-specific number of threads for parallel computation (`-t`), names of folders containing output of previous analysis steps, or parameters left at their default values. nb. = number; frac. = fraction; avg. = average; aln. = alignment; min. = minimum; max. = maximum.

| Step(s) | Var | Parameter | Iteration 1 | Iteration 2 |
|---|---|---|---|---|
| 1+2+3+6 | $s | sample file | samples.dalbergia.12.txt | samples.dalbergia.txt |
| 1+3+4 | $1 | reference sequences | Cajanus_cajan_6555reg.fasta | Dalbergia_iter1_3736reg.fasta |
| 1 | $T | bwa-mem alignment score | 10 | 10 |
| 1+2 | $Q | min. mapping quality | 10 | 10 |
| 1 | $e | trimmed read file extension | .trim1.fastq.gz,.trim2.fastq.gz | .trim1.fastq.gz,.trim2.fastq.gz |
| 1+4+5+7 | $2 | sample / taxon group file | mapfile.dalbergia.12.txt | mapfile.dalbergia.txt |
| 1 | $3 | min. frac. regions | 0.2 | 0.2 |
| 1 | $4 | min. frac. taxa | 0.4 | 0.7 |
| 1+3 | $5 | min. aln. length | 1 | 1 |
| 1 | $6 | min. avg. coverage | 8 | 8 |
| 1 | $7 | max. avg. coverage | 1000 | 1000 |
| 1 | $8 | min. aln. fraction | 0 | 0 |
| 1 | $9 | min. frac. conforms | 0.4 | 0.7 |
| 3 | $10 | min. normalized score | 2 | 2 |
| 4+5 | $11 | min. frac. regions | 0.2 | 0.7 |
| 4+5 | $12 | min. frac. taxa | 0.5 | 0.85 |
| 4+5 | $13 | max. nb. contigs | 2 | 2 |
| 4+5 | $14 | min. normalized score | 2 | 2 |
| 4+5 | $15 | min. aln. length | 80 | 80 |
| 4 | $16 | min. aln. fraction | 0 | 0 |
| 4 | $17 | min. score | 1 | 1 |
| 4 | $18 | min. contig length | 1 | 1 |
| 4+5 | $19 | min. frac. conforms | 0.5 | 0.7 |
| 6 | $20 | min. merging score | 0.9 | 0.95 |
| - | $21 | max. gap ratio | - | 0.3 |
| - | $22 | max. nucleotide diversity | - | 0.15 |

*Step 1: Read mapping*

We ran *BWA* version 0.7.12-r1039 (Li & Durbin, 2009) and *BWA-MEM* in the `$mappingdir` directory, using the quality-trimmed and quality-filtered reads with the indicated file extensions (-e option, comma-separated string denoting file extensions of forward and reverse reads, respectively) located in the `$trimmedreads` directory (-d option), and the respective reference sequences for each taxon set and iteration (-r option). The script outputs reads with a minimum alignment score of `$T` (-T option), marks secondary hits, and only retains reads with a minimum mapping quality of `$Q` (-Q option) in the final SAM files before compressing them to BAM format.

An early filtering of target regions with inadequate coverage across samples prevents time-consuming sequence assembly of target regions that would likely be filtered out in step 4. Computations were performed for all samples specified in `$s` (-s option) using 4 times 5 threads in parallel (-t option) as follows:

```
run.bwamem.sh -s $s -r $1 -e $e -T $T -Q $Q -d $trimmedreads -t 4
```

We performed coverage analysis on the BAM files filtered for mapping quality equal or above `$Q` (-Q option) for each sample, and wrote all coverage results to one file as follows:

```
get.coverage.stats.sh -s $s -Q $Q -t 20
collect.coverage.stats.R $s $Q
```

We implemented seven filtering criteria to identify target regions with adequate average coverage across the taxon groups specified in `$2`. The first two filters take absolute thresholds and aim to remove poorly sequenced samples or target regions: `$3` minimum fraction of regions with at least one mapped read in a sample (filters samples), `$4` minimum fraction of samples with at least one mapped read in a region (filters target regions). The next four filters take thresholds that need to be met in a specified fraction of samples in each considered taxon group: `$5` minimum BWA-MEM alignment length, `$6` minimum average coverage in the aligned region, `$7` maximum average coverage in the aligned region, `$8` minimum alignment fraction (BWA-MEM alignment length divided by target region length). `$9` is the minimum fraction of samples in each taxon group that need to pass each filter in order to keep a certain target region.

```
filter.visual.coverages.R $2 coverage_stats.txt $1 $3 $4 $5 $6 $7 $8 $9
```

This script visualized the coverage statistics as violin plots and heatmaps, and saved a list of kept samples (`$s`) as well as a list of kept regions (`$1`) for sequence assembly.

*Step 2: Sequence assembly*

We extracted read pairs from quality-filtered and quality-trimmed reads located in the `$trimmedreads` directory (`–d` option). The `-s` and `-l` parameters are used to pass the list of samples and loci (target regions) to be processed in parallel, respectively. This step was carried out on a local scratch (`$extractedreads` directory) using 20 parallel threads (`–t` option). At least one of the two reads per extracted read pair mapped to a retained target region with a minimum mapping quality of 10 (`–Q` option):

```
extract.readpairs.sh -s $s -l $1 -d $trimmedreads -m $mappingdir -Q $Q -t 20
```

We assembled the extracted reads located in the `$extractedreads` directory (`–r` option) into consensus contigs (contigs hereafter) separately for each sample and retained region using *dipSPAdes* (*SPAdes* version 3.6.0) in '`assembly-only`' and '`careful`' mode, with an automatic coverage cutoff. This step was carried out on a local scratch (`$assemblies` directory) using 20 parallel threads (`–t` option):

```
run.dipspades.sh -s $s -r $extractedreads -t 20
```

*Step 3: Orthology assessment*

We ran EXONERATE version 2.2 for each sample and each retained target region (`-l` option) with the '`affine:local`' and '`exhaustive`' options, using the contigs located in the `$assemblies` directory (-d option) as query sequences and the target regions (`-r` option) as target sequences. We stored alignment statistics of all consensus contigs that aligned to the same target region in the `$exonerate` directory (`-d` option), but limited the report to the best alignment per contig as follows:

```
select.best.contigs.per.locus.sh -s $s -l $l -r $1 -d $assemblies -t 20
```

Contigs with a target alignment length of at least the specified threshold (`-a` option) and a normalized alignment score (defined as the raw EXONERATE alignment score divided by the target alignment length) of at least the specified threshold (`-c` option) were considered as potentially homologous and retained. If more than one contig met these requirements, and if none of these contigs physically overlapped based on the alignment statistics, the best-matching contig was combined with the additional contig(s) using an appropriate spacer and by taking the directionality into account as follows:

```
combine.contigs.parallel.sh -s $s -d $exonerate -a $5 -c $10 -t 20
```

We collected the EXONERATE statistics of each sample and plotted the number of contigs per target region for the different taxon groups as follows:

```
collect.exonerate.stats.R $s $exonerate
plot.contig.numbers.R regions_contignumbers.txt $2
```

*Step 4: sample and region filtering*

We implemented nine filtering criteria to identify target regions with adequate assembly quality across the taxon groups specified in `$2`. The first two filters take absolute thresholds and aim to remove poorly assembled samples or target regions: `$11` minimum fraction of regions with at least one contig in a sample (filters samples), `$12` minimum fraction of samples with at least one contig in a region (filters target regions). The next five filters take thresholds that need to be met in a specified fraction of samples in each considered taxon group: `$13` maximum number of non-zero (fragments combined) contigs in a target region, `$14` minimum normalized EXONERATE alignment score, `$15` minimum EXONERATE alignment length, `$16` minimum alignment fraction (EXONERATE alignment length divided by target region length), `$17` minimum raw EXONERATE alignment score, `$18` minimum contig length. `$19` is the minimum fraction of samples in each taxon group that need to pass each filter in order to keep a certain target region.

```
filter.visual.assemblies.R $2 loci_stats.txt $1 $11 $12 $13 $14 $15 $16 $17
$18 $19
```

*Step 5: Target region alignment and alignment trimming*

We generated multifasta files in the `$multifasta` directory for all retained target regions, containing all retained contigs and samples as follows:

```
taxa=taxa_kept-$11.txt
regions=regions_kept-$11-$12-$13-$14-$15-$19.txt
create.multifastas.parallel.sh -s $taxa -l $l -d $exonerate -t 20
```

We generated alignments in the `$mafft` directory using MAFFT version 7.123b, the '`localpair`' and '`adjustdirection`' flags, and 1000 maximum iterations:

```
align.multifastas.parallel.sh -d $multifasta -m 'localpair' -t 20
```

Raw alignments were trimmed at both ends until an alignment site had nucleotides in at least 50% of aligned sequences (`-c` option) and a maximum nucleotide diversity (i.e., the sum of the number of base differences between sequence pairs divided by the number of comparisons) of 0.25 (`-n` option). The `-v` flag triggered visualization of the alignment end trimming procedure. Trimmed alignments were written to the `$endtrimmed` directory as follows:

```
trim.alignment.ends.parallel.sh -s $2 -d $mafft -c 0.5 -n 0.25 -t 20 -v
```

Internal trimming was carried out by first removing any alignment site with nucleotides in less than 40% of aligned sequences (`-c` option). Potential mis-assemblies or mis-alignments in each sequence were resolved using a sliding window approach with window size 20 (`-z` option) and step size 1 (`-S` option). Specifically, we trimmed windows at contig ends if more than 50% of the nucleotides in the conserved part of the window deviated from the alignment consensus (`-n` option). The script defines a conserved part of each window as the alignment sites with nucleotides in at least 20% of samples, and where the frequencies of minor alleles are all below 30% without considering gaps. After window-based trimming, the script also removes sites with sequence data for less than the specified fraction of aligned sequences (`-c` option) again. The `-v` flag triggered visualization of the internal trimming procedure with sliding window approach. Trimmed alignments were written to the `$trimmed` directory as follows:

```
trim.alignments.parallel.sh -s $2 -d $endtrimmed -c 0.4 -z 20 -n 0.5 -S 1 -t 20
-v
```

*Step 6: Merge overlapping alignments*

We calculated a consensus sequence for each end-trimmed and internally trimmed alignment located in the directory `$trimmed`, using a minimum allele frequency of 1 (`-m` option) to call IUPAC ambiguity and a minimum base frequency of 0.01 (`-b` option) to return a consensus instead of a gap. This parameter combination ensured that the most frequent allele was called at each alignment site rather than IUPAC ambiguity codes or gaps. If two alleles were equally frequent at any alignment site, one was randomly sampled to represent the consensus. The `-g` flag ensured that gaps were removed from the final consensus sequence, and the -n flag ensured that completely ambiguous consensus bases (Ns) were removed from the final consensus sequence. The `-v` flag triggered visualization of the consensus calculation:

```
get.consensus.from.alignment.parallel.sh -s $taxa -d $trimmed -m 1 -b 0.01 -t
20 -gnv
```

We renamed the sequence names of alignment consensus sequences stored in `$cons` to dispose of the suffix added during alignment and trimming before identifying the best non-reciprocal BLAST+ hits between alignment consensus sequences as follows:

```
rename.fasta.headers.R $cons ".all.aln.etr.itr" FALSE FALSE
blast.vs.self.sh $cons
```

We then identified BLAST+ hits at alignment ends and stored a list of physically overlapping alignments (names of overlapping target regions on the same line) as follows:

```
cbase=$(basename $cons .fasta)
find.overlapping.alignments.R $cbase.vs.self.blast.filtered TRUE 'LG_' '_'
```

Arguments 2–4 limit the identification of overlapping alignments to target regions of the same linkage group, using specific strings surrounding a linkage group identifier present in target region names. We then aligned all contigs of up to five physically overlapping target regions using the same alignment algorithm as before. Merged alignments were written to the `$merged` directory as follows (visualization was triggered by default):

```
overlaps=$cbase.list
align.overlapping.contigs.sh -l $overlaps -c $multifasta -m 'localpair' -t 20
```

In cases where contigs of the same sample overlapped with a mismatch, only the base with higher frequency at that alignment site was considered. A success score of each merging procedure was computed based on the number of mismatches in overlapping contigs of the same individuals relative to the total number of bases in the alignment. The success score amounted to 1 if there were no mismatches in any individual. We discarded any merged alignment with a score smaller than 0.85 (subfamily set) or 0.9 (species set).

```
filter.merged.alignments.sh -d $merged -s $20
```

Unsuccessfully merged alignment sets were visually inspected to identify whether some subsets of alignments sufficiently overlapped to allow for merging. The manually selected alignments were merged again and combined with the automatically merged alignments if they showed a sufficient success score. All successfully merged alignments were then trimmed as before and used as replacements for overlapping alignments.

```
trim.alignment.ends.parallel.sh -s $s -d $merged -c 0.5 -n 0.25 -t 20 -v
trim.alignments.parallel.sh -s $s -d $merged -c 0.4 -z 20 -n 0.5 -S 1 -t 20 -v
replace.overlapping.alignments.R $trimmed $merged $overlaps
```

*Step 7: Create representative reference sequences*

Sets of reference consensus sequences for different taxon groups were generated, combined, aligned, and a group consensus was derived as follows:

```
get.group.consensus.sh -s $2 -d $trimmed -m 1 -b 0.01 -z ".all.aln.etr.itr.cons"
-t 20 -gnv
```

The resulting FASTA file `$newref` was renamed according to the taxon set and number of remaining target regions:

```
rename.fasta.headers.R $newref ".cons.aln" FALSE FALSE
mv $newref <SET NAME>_iter<# ITERATION>_<# REGIONS>reg.fasta
```

5.3.6 | Phylogenetic analyses

Supermatrix (concatenation) approach

We assessed alignments as follows:

```
assess.alignments.parallel.sh -f $trimmed -t 20
```

We filtered out alignments with excessive gap ratio or nucleotide diversity as follows:

```
filter.visual.alignments.R $trimmed.assess.txt $21 0 $22 100
```

We concatenated alignments as follows:

```
concatenate.fastas.R $trimmed
```

We ran maximum likelihood search on the concatenated alignments using RAxML as follows:

```
raxmlHPC-PTHREADS-SSE3 -f a -m GTRCAT -x 85397 -p 24686 -s $trimmed.phy
-n $trimmed.catBS100.nex -T 20 -N 100 > catBS100.log 2> catBS100.err
```

Gene tree summary approach

We generated gene trees for different alignments as follows:

```
get.gene.trees.parallel.sh -d $trimmed -n 100 -t 20
```

We collapsed branches with low support (below 10) as follows:

```
nw_ed $trimmed.genetrees 'i & b<=10' o > $trimmed.BS10.genetrees
```

We inferred the species tree using ASTRAL-III as follows:

```
java -Xmx1G -jar astral.5.6.3.jar -i $trimmed.BS10.genetrees -o $trimmed.BS10.single.spectree -t 2 2> $trimmed.BS10.single.log
```

We added 0.5 coalescent units to each zero terminal branch length as follows:

```
add.to.terminal.branches.R $trimmed.BS10.single.spectree 0.5
```

5.3.7 | Popluation genetics analyses

We removed PCR duplicates and capped BAM files as follows:

```
bsub < remove.dups.and.cap.lsf
```

We collected a table of percentage of PCR duplicates as follows:

```
grep '^LIBRARY' -A1 ${folder}/stats_dup/*txt --no-group | awk '!seen[$3]++' | cut -f1,9 | cut -f3 -d'/' | sed -e 's/.dupstats.txt-Unknown Library//' > percdup.txt
```

We called SNPs on the EULER cluster as follows:

```
module load gcc/4.8.2 gdc python/2.7.11 perl/5.18.4 samtools/1.3

folder="Chapter1.3_mapsnp-2396_51.nodup.cov500"
ref="Dalbergia_4c2i_2396reg.fasta"
maxlen=10000
lenperjob=10000

## Split jobs by regions
samtools faidx ${ref}
fasta_generate_regions.py ${ref} ${maxlen} > regions.txt
split.freebayes.regions.file.pl regions.txt ${lenperjob}
mkdir regions
mv regions_${lenperjob}_* regions

## Create output dir
mkdir vcfs
ls -1 ${folder}/*.bam > bamlist.txt

## Submit job
bsub < submit.multi.freebayes.cmds.lsf

## Combine .vcf files
combine.vcf.files.sh vcfs

## Rename single .vcf file
mv vcfs.vcf Dalbergia_51_2396_203916.vcf
```

We filtered raw variants as follows:

```
filter.snps.sh -v Dalbergia_51_2396_203916.vcf -r $ref -n Dalbergia_51_2396_
```

The above script uses *vcftools* to filter for mapping quality (`--minQ 30`), minimum depth (`--minDP 3`), and minimum mean depth (`--min-meanDP 10`), decomposes complex variants using *vcflib*'s *vcfallelicprimitives* functionality, and finally removes insertions and deletions (`--remove-indels`).

We created a genind object from the filtered SNPs for analyses using the R ADEGENET package as follows:

```
vcf=Dalbergia_51_2396_116500.filtered.vcf
grep '^>' $ref | cut -f2 -d'>' > regions
subset.vcf.by.region.sh -v $vcf -r regions -t 20
vcf2adegenet.R $(basename $vcf .vcf) $ref 1> get_gi.log 2> get_gi.err
```
